# Evolution Tracks Multiple Timescales at Once

**DOI:** 10.1101/2022.10.27.514123

**Authors:** Emily L. Behrman, Paul Schmidt

## Abstract

Adaptation is often modeled as an immediate response to the environment currently experienced, yet we demonstrate that past conditions can continue to shape future genomic change, independent of the present environment. Using seven years of longitudinal genomic, environmental, and phenotypic data from a wild *Drosophila melanogaster* population, spanning more than 75 generations, we identify two genomically independent temporal modes of adaptation: a concurrent mode, tracking the current environment, and a lagged mode, carrying the imprint of a prior period forward. Despite acting on largely distinct genetic variants, both modes converge on shared neural and developmental pathways, most strongly in genes with broad phenotypic effects, suggesting that the circuitry linking environmental sensing to behavior is a recurring target of multi-temporal selection. These results show that adaptation can proceed simultaneously across distinct timescales, a pattern that may be widespread wherever selection recurs in structured, predictable ways.

## Main text

How populations adapt to environmental change is a central question in evolutionary biology. Most adaptation is thought to be polygenic, yet our understanding of selection dynamics comes largely from traits with simple genetic architectures (*1*). This gap is compounded by the temporal variability of most natural environments, a condition most models of adaptation do not address: fitness optima shift, trade-offs reverse, and the axes of selection change from one generation to the next (*2*, *3*). Classic studies of selection following extreme climatic events show that the severity of a selective episode shapes the strength and direction of the evolutionary response (*3*, *4*), but whether such consequences propagate to structure future evolution remains largely untested. Resolving this requires longitudinal integration of environmental drivers, genomic responses, and phenotypic outcomes within a single natural population (*5*).

Longitudinal sampling turns environmental variability from a confounding factor into a diagnostic strength: tracking real year-to-year differences reveals which environmental features and temporal dynamics actually drive adaptation. Existing genome-environment association (GEA) approaches, however, generally assume selection is immediate (*6*). This leaves potential carry-over effects untested, despite being established at the phenotypic level (*7*), and rarely incorporates phenotype itself, the trait-level outcome that genomic and environmental change are expected to produce.

Seasonally fluctuating environments offer a tractable system for testing whether adaptation is temporally partitioned. *Drosophila melanogaster* persists year-round in temperate regions, overwintering in a dormant state and emerging synchronously each spring (*8*, *9*), with winter acting as the dominant selective force of the annual cycle. Existing genomic studies of seasonal adaptation in this system have largely treated winter and summer selection as a fixed, year-independent contrast (*10–12*), overlooking whether selection at one point structures the next. Field-based experimental evolution has tested specific within-season hypotheses (*13*, *14*), but provides only a snapshot; it has not been paired with the longitudinal measurements needed to ask whether selection in one period structures the evolutionary trajectory of the next.

We combined seven years of common-garden phenotyping with longitudinal pooled whole-genome sequencing from a single *D. melanogaster* population (Fig. 1), fitting allele-frequency dynamics to multivariate environmental drivers across multiple temporal scales. This reveals two genomically independent temporal modes of adaptation—one tracking the current environment, the other carrying the imprint of a prior period forward—that converge on shared neural and developmental pathways despite acting on largely distinct genetic variants.

**Fig. 1.**
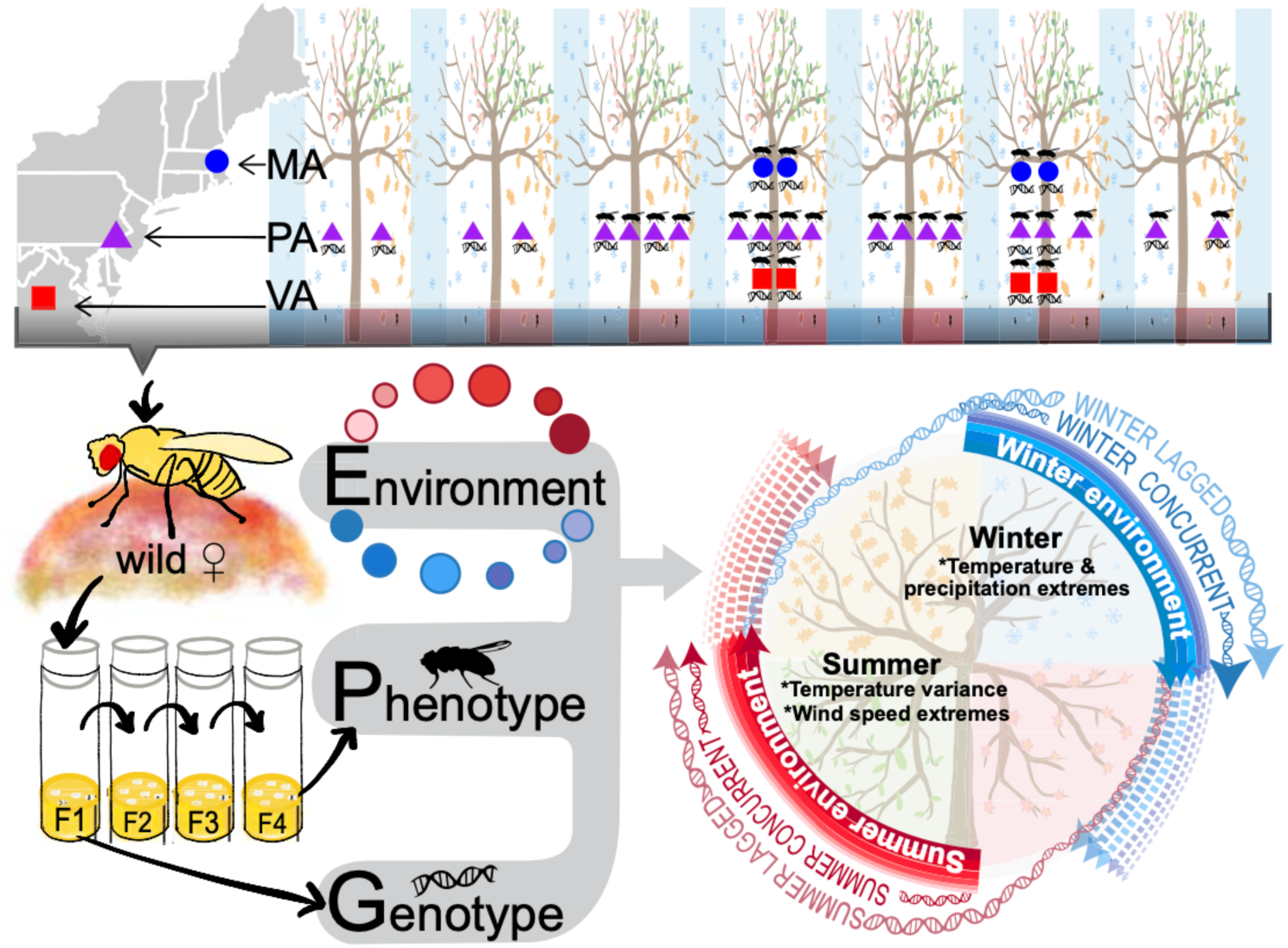
Longitudinal sampling design and two independent temporal modes of seasonal adaptation. Wild-caught females were collected at multiple seasonal timepoints (June-November) across seven years (2009–2015) from three mid-Atlantic populations spanning a latitudinal cline: Massachusetts (MA; 42.5°N), Pennsylvania (PA; 39.9°N), and Virginia (VA; 38.0°N). Fly icons indicate phenotypic sampling; DNA icons indicate genomic sampling. At timepoints with both icons, each wild-caught female founded an isofemale line maintained in a common laboratory environment for four generations, with genotype (G) characterized by pooled genome sequencing (pool-seq) of F1 males and phenotype (P) assayed on F4 females; genotype-only timepoints were pool-sequenced directly from wild-caught flies without establishing isofemale lines. Seasonal environmental (E) variation was summarized using six principal components, a statistical technique that condenses many correlated meteorological measures (temperature, precipitation, snow, and wind) into a smaller number of composite variables, together explaining 99.4% of environmental variance across the measures. Allele frequency changes were correlated with environmental composite scores to identify two independent temporal modes of seasonal adaptation: a concurrent mode in which winter and summer allele frequency changes respond to the current season’s environment, and a lagged mode in which one season’s environment drives allele frequency change in the following season.

### Phenotypic tracking is driven by seasonal bottlenecks and partitioned plasticity

Phenotypes changed rapidly and repeatedly within seasons across multiple years for all four traits assayed (development time, starvation resistance, heat knockdown, and chill recovery), with significant changes between temporally adjacent collections (Fig. 2A). This seasonal change was validated by parallel phenotypic differentiation across space: our focal population sits at the center of a latitudinal transect spanning at least 4° in the mid-Atlantic United States, flanked by populations equidistant to the north and south (Fig. 2A). Populations across this transect also evolved across seasons, and for two traits, development time and starvation resistance, the seasonal signal actually exceeded the spatial one (χ2=144 vs 24 and χ2=7.4 vs 5.2, respectively). Winter severity was the primary determinant of the magnitude of change occurring the subsequent summer: rapid phenotypic evolution unfolded against a backdrop of 11–18 estimated non-overlapping generations per growing season from spring to autumn (fig. S1A), with harsh winters acting as a strong, discrete selective bottleneck that narrowed the population to its stress-resistant winter survivors (Fig. 2B, fig. S2). The subsequent summer represents a release from that bottleneck as populations expand under environmental conditions favorable for growth; mild winters fail to impose this bottleneck, and the following summer’s phenotypic trajectories are correspondingly undirected.

**Fig. 2.**
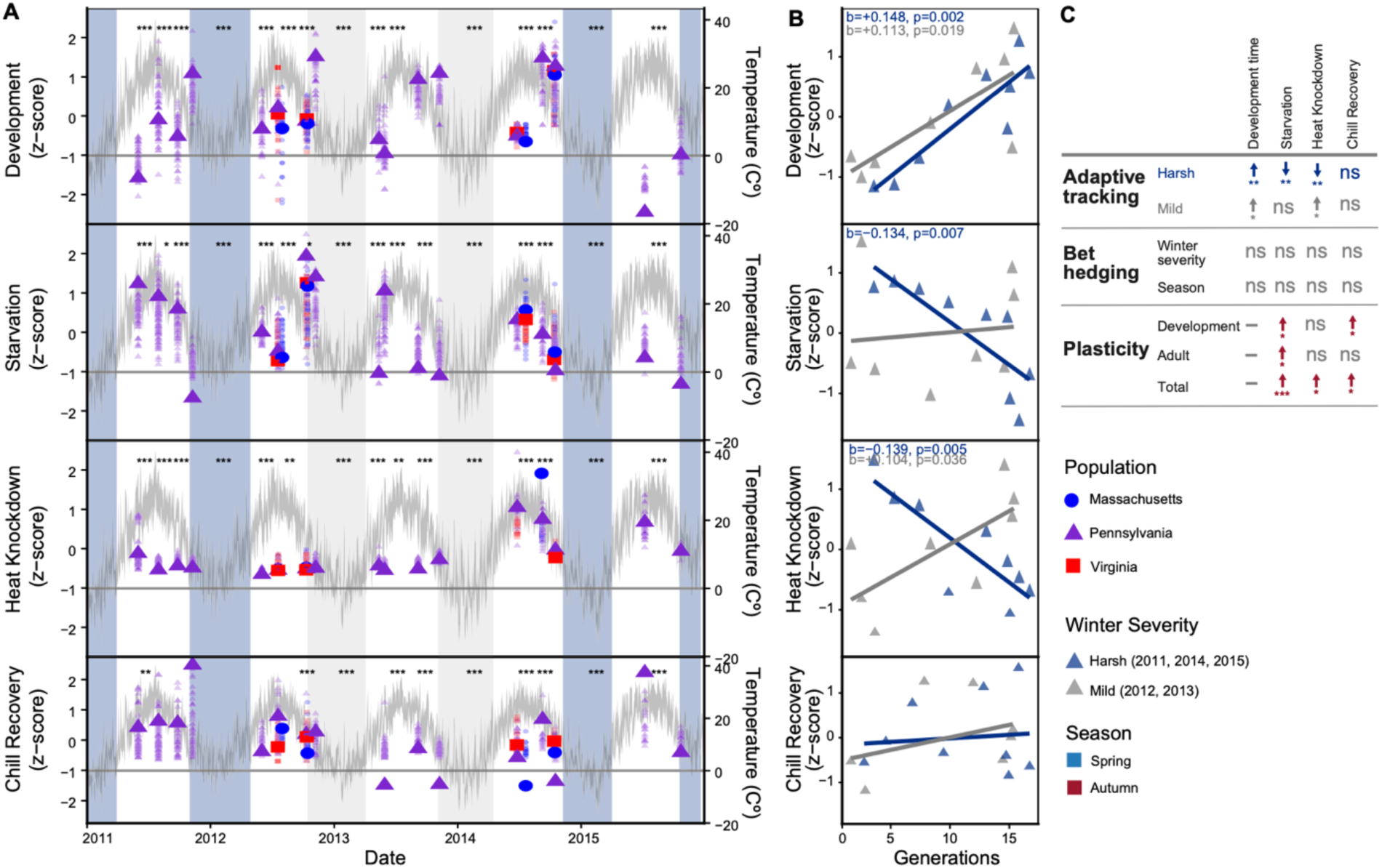
Seasonal phenotypic adaptation is genetically based and environmentally structured. **(A)** Phenotypic z-scores for development time, starvation resistance, heat knockdown, and chill recovery, Massachusetts (blue circles), Pennsylvania (purple triangles), and Virginia (red squares), 2011-2015. Gray line, daily temperature at the Pennsylvania site; blue/gray shading, harsh/mild winters. Asterisks, significant differences between adjacent Pennsylvania collections (Wilcoxon test, BH-corrected; *P<0.05, **P<0.01, ***P<0.001). (B) Standardized trait values versus cumulative generations since dormancy, Pennsylvania, fit separately for harsh (blue) and mild (gray) winters (Methods); shown for significant fits only. **(C)** Summary of phenotypic response modes (full data, fig. S2). Adaptive tracking, direction/significance of the trend in B. Bet-hedging, variability among a line’s replicate measurements within one collection (coefficient of variation), compared between winter severities or seasons. Plasticity, breadth of the trait’s response across different rearing temperatures, spring versus autumn. Arrows, direction of significant effects (blue, harsh/spring; red, autumn; gray, mild); ns, not significant. BH-corrected throughout.

Winter severity was the primary determinant of the predictability and magnitude of phenotypic change. Development time tracked cumulative generations consistently regardless of winter severity, a reliably predictable response independent of context. Starvation resistance tracked only after harsh winters, with no significant relationship after mild winters: its predictability, not just its presence, is contingent on the strength of winter selection. Heat knockdown tracked generations regardless of severity but with opposite slopes (Fig. 2B), showing that severity determines not just whether the response is predictable but which direction it runs. These three distinct architectures show that winter severity shapes phenotypic predictability as much as magnitude and are explored further in the following genomic analyses.

Phenotypic plasticity operates as a second, temporally partitioned strategy alongside the phenotypic tracking described above. Total plasticity assessed across developmental and adult reaction norms was greater in autumn than spring for starvation resistance, chill recovery, and heat knockdown (Fig. 2C, fig. S1B), suggesting summer selection favors flexible responsiveness within a season, though this was assessed in a single year and would benefit from replication. Bet-hedging, measured as within-collection variance among a line’s replicate measurements, showed no seasonal or severity-dependent pattern for any trait (Fig. 2C, fig. S1C). Together, we show temporally partitioned components of a single strategy for coping with environmental heterogeneity where plasticity manages within-season uncertainty and phenotypic tracking provides the directional signal that connects seasons across years.

### Concurrent genomic responses are decoupled across seasonal environmental axes

Winter and summer selection drove qualitatively different genomic responses in both scale and consistency. We identified concurrent SNPs whose allele-frequency trajectory over summer or winter during the seven years correlated with that season’s value for at least one of six environmental principal components, which together explained 99.4% of environmental variance (Fig. 3A, fig. S3A). This approach identified 65,403 unique concurrent GEA SNPs meeting jackknife-based significance criteria, of which only 1,229 were significant in both seasons (Fig. 3B, fig. S3, B and C). Winter- and summer-associated SNPs were largely non-overlapping and in linkage equilibrium with each other, a pattern not explained by chromosome- or inversion-level enrichment (fig. S3, D to F).

**Fig. 3.**
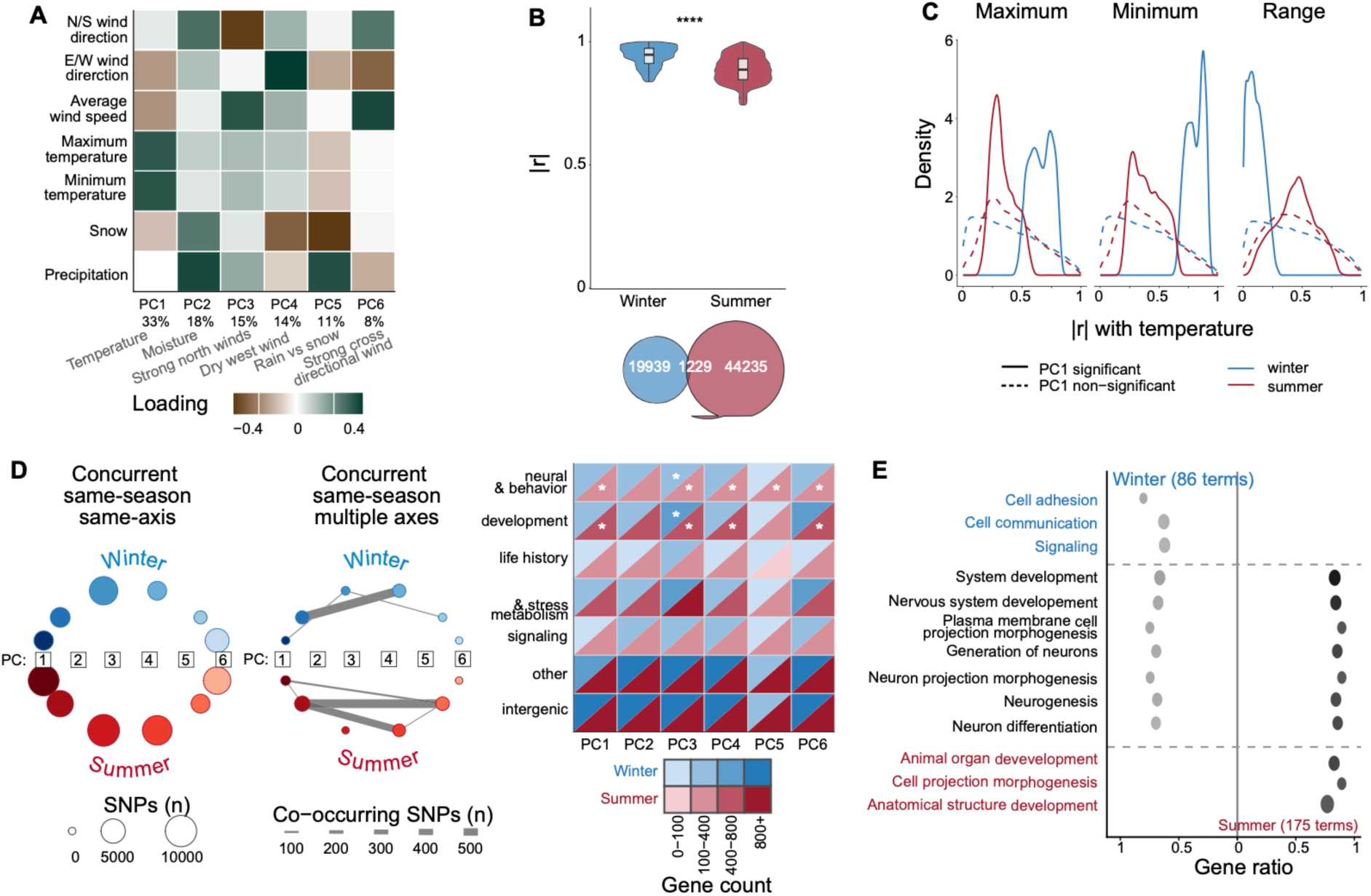
Multivariate seasonal environments drive neural and behavioral genomic responses with greater predictability under winter selection. **(A)** The contribution of each raw weather variable to a given principal component (PC) at the Pennsylvania site; six PCs explain 99.4% of variance (shown below). **(B)** Jackknife-corrected |r| for winter-versus summer-associated SNPs (Wilcoxon test, BH-corrected; **“P<0.0001); Venn, SNPs significant in winter only, summer only, or both (n=1,229 shared). **(C)** Density of |r| with temperature (maximum, minimum, range), PC1-significant (solid) versus non-significant (dashed) SNPs, winter (blue) and summer (red). **(D)** Left, winter-concurrent (top) and summer-concurrent (bottom) PCs shown as nodes, sized by the number of SNPs significant for that PC. Right, gene counts by functional category and PC for SNPs unique to one season, shown as diagonal heatmap with winter (upper-left) and summer (lower-right) triangles per cell, fill, gene count; stars, categories independently enriched for that PC and season. **(E)** GO enrichment, winter-only (blue font) versus summer-only (red font) and shared (gray font) SNPs (BH-corrected P≤0.05).

Winter selection produced a concentrated genomic signal, while summer produced a broader but weaker signal, with roughly twice as many SNPs (21,168 vs. 45,464) but lower correlation strength (mean |r|=0.940 vs. 0.887; Wilcoxon p<2.2x10-16; Fig. 3B, fig. S3G), reflecting greater predictability of the genomic response to winter’s environment. This advantage held for individual PCs associated with temperature, moisture, and precipitation after accounting for winter’s smaller number of seasonal transitions, but not for the wind-related axes, where summer matched or slightly exceeded winter (fig. S3H). Winter- and summer-significant SNPs also tracked different features of the same variable: temperature extremes for winter, temperature range for summer (Fig. 3C).

Concurrent selection operates along largely independent environmental axes. Most SNPs were specific to a single season and a single environmental PC, with only ∼3% of season-specific SNPs tracking more than one PC within or across seasons (Fig. 3D, fig. S3I). Where within-season, multi-axis connectivity did occur, winter’s was more concentrated than summer’s, forming one tightly coupled hub (moisture, PC2; dry west wind, PC4), while summer’s connections were more diffuse. Cross-season connectivity at the same PC also exhibited structure of winter’s influence propagating into a broader, more diffuse summer response, rather than the two seasons behaving as independent or symmetric regimes (fig. S3I).

### Identification of a lagged genomic mode independent of current selection

We identify a second, largely independent mode of lagged genomic response, where allele-frequency change in one season correlates with the environment of the season before it (lagged GEA, fig. S4A), distinct from the current season’s own environment (concurrent GEA, fig. S3A). Winter-lagged SNPs reflect the summer environment predicting winter allele-frequency change, and summer-lagged SNPs the reverse. Testing both directions identified 16,391 significant lagged SNPs that were largely distinct from the concurrent GEA SNPs; only 8.44% overlapped between the two (Fig. 4A, fig. S4, B and C). As with the concurrent signal, lagged PC-level associations reflected environmental variation: lagged SNPs showed higher |r| with the corresponding raw environmental variables than non-significant SNPs, across all six PCs (fig. S5). Two separate checks confirmed that concurrent and lagged signals are genuinely independent, not just two readouts of the same physically linked variants. First, even where a lagged and concurrent SNP sit close together, they are rarely in strong linkage (median haplotype r2=0.267 at <50bp; Fig. 4B). Second, physical proximity between the two signals is itself uncommon; lagged SNPs were enriched for a nearby concurrent SNP only within 25bp, with no such enrichment beyond 50bp (Fig. 4C).

**Fig. 4.**
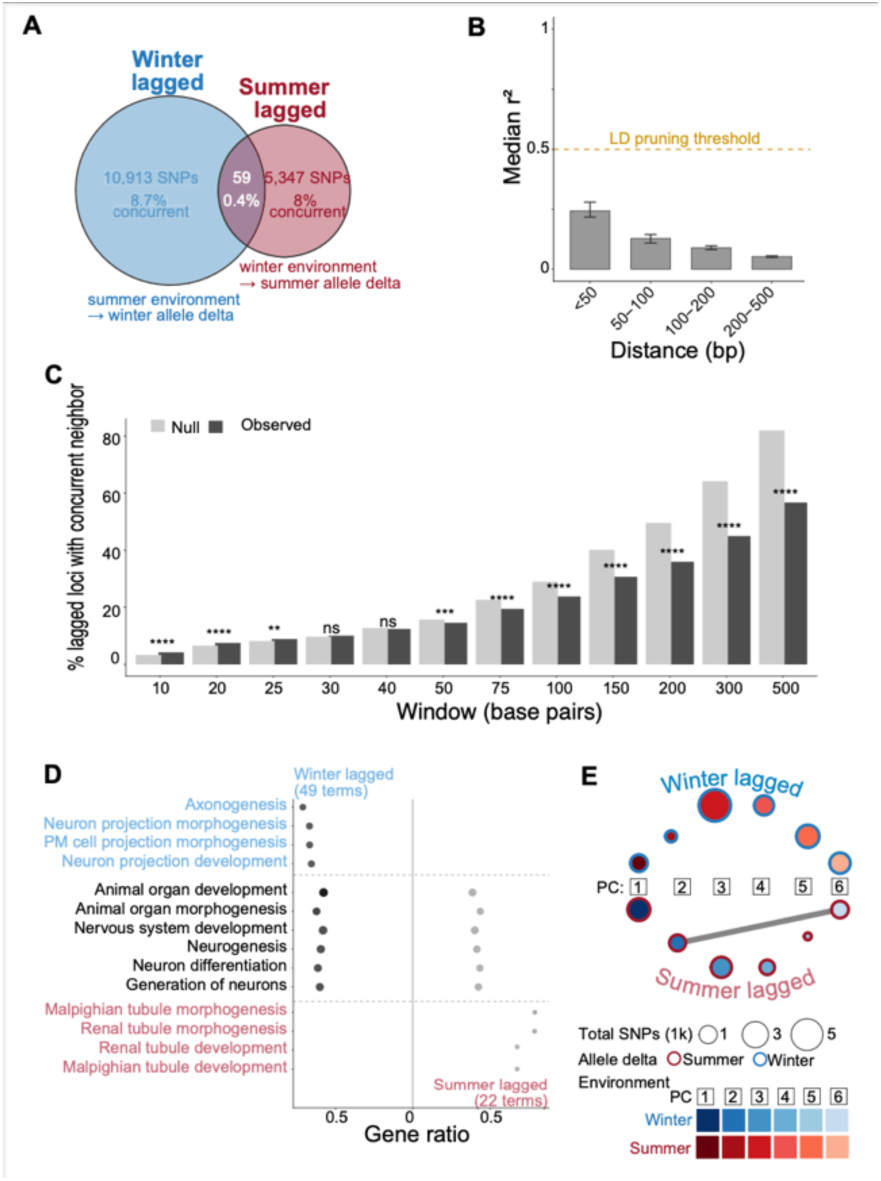
Lagged genomic adaptation is asymmetric and largely independent of concurrent responses. **(A)** Venn, SNPs significant winter-lagged only, summer-lagged only, or both (n=59 shared); labels, total SNPs per direction and percent overlap with concurrent GEA. **(B)** Median haplotype r2 measures how often two genetic variants are inherited together between physically proximate SNP pairs, by distance bin; dashed line, linkage disequilibrium (LD)-pruning threshold (r2=0.5), the cutoff above which nearby SNPs are treated as redundant. **(C)** Percent of lagged SNPs with a nearby concurrent SNP, by window size, compared against a null expectation; enrichment at ≤25bp and depletion beyond 50bp indicate that physical proximity between the two signals is uncommon beyond a narrow window. **(D)** GO enrichment, winter-lagged (blue font) versus summer-lagged (red font) and shared (gray font) SNPs, by functional category. **(E)** PC-routing web showing how often SNPs for different PCs co-occur within the same gene, for winter-lagged and summer-lagged SNPs, with per-PC total-SNP heatmap; edge width is shared-SNP count; node outline color is allele-frequency delta season.

The lagged signal was asymmetric in count between directions, though not in predictability. Winter-lagged SNPs outnumbered summer-lagged SNPs more than two to one (10,913 vs. 5,347; Fig. 4A), with only 59 SNPs significant in both. This signal was not concentrated in a small number of overactive loci, a few chromosomal regions, or cosmopolitan inversions, but was instead driven by many SNPs each tied to a single environmental PC (fig. S4E).

### Genomic responses are organized by environmental trigger rather than temporal mode

SNPs were categorized by two independent features: which season’s environment triggered the response, and in which season the resulting allele-frequency change occurred (Fig. 5A). Winter environment shapes the allele-frequency change of both winter-concurrent and summer-lagged SNPs; the same holds in reverse for summer. This framework lets us ask whether genomic responses are organized primarily by temporal mode (concurrent versus lagged) or independent of mode by environmental trigger season (winter versus summer).

**Fig. 5.**
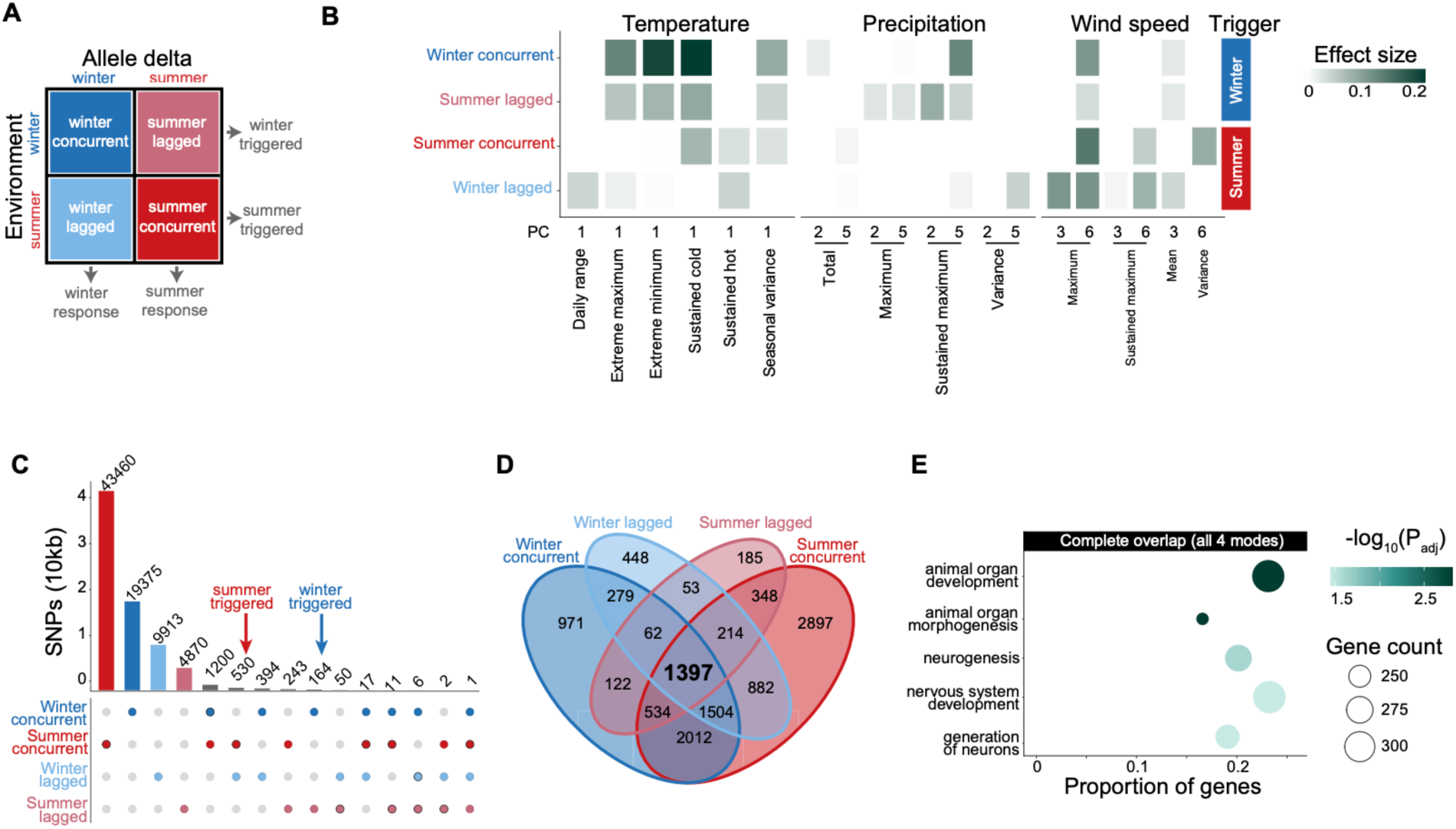
Concurrent and lagged responses converge on a shared set of functionally coherent genes, despite genomic independence. **(A)** Conceptual framework: environment and allele-delta timing define concurrent/lagged categories; winter-/summer-triggered groupings indicated. **(B)** Winter-/summer-triggered environmental signature by variable (temperature, precipitation, wind speed). **(C)** SNP overlap among all combinations of the four categories; winter-/summer-triggered shared SNPs indicated. **(D)** Four-way Venn, genes hit by winter-concurrent, summer-concurrent, winter-lagged, and summer-lagged SNPs; complete overlap (n=1,397) is smaller than expected from permutation of matched random SNPs sets. **(E)** GO enrichment, 1,397 complete-overlap genes; no significant terms for winter-only, summer-only, or partial-overlap gene sets.

The environmental features associated with genomic change differed by trigger season, not by temporal mode, reflecting that winter and summer are simply different selective forces (Fig. 5B, fig. S5). Winter-triggered categories showed a broad temperature signature, particularly shaped by temperature extremes, with a secondary precipitation signal that concentrated most clearly in summer-lagged SNPs. Summer-triggered categories showed a different temperature signature centered on temperature variation and a substantial but non-specific role for wind speed (Supplementary Materials). Together, these patterns show that winter and summer selection are not simply weaker or stronger versions of each other, but respond to genuinely different features of the environment.

The shared organization by trigger extended to gene identity, though rarely to individual SNPs (Fig. 5C), consistent with the genetic redundancy expected under polygenic adaptation (*15*). We identified a set of 1,397 genes containing SNPs hit by all combinations of modes and triggers (Fig. 5D). This complete overlap was significantly smaller than expected under two independently constructed permutation nulls, with the observed count falling below every one of 200 permutations in both (empirical P<0.005). This is consistent with these four selective processes being largely independent of one another. We used the overlap genes for downstream functional enrichment testing to see if the different selection regimes converge on shared biological functions.

### Multi-temporal selection converges on conserved neural and developmental circuitry

Despite responding to different environmental features, neural and developmental function was the recurring target of seasonal genomic change. Gene ontology (GO) terms shared between winter and summer for concurrent GEA were dominated by neural and developmental processes, while winter’s distinct terms centered on cell communication and signaling, and summer’s on organ and anatomical morphogenesis (Fig. 3E). At the level of individual PCs, neural and developmental function were the only categories to reach independent enrichment (Fig. 3D). Lagged GEA showed similar convergence, with shared terms spanning broader organ and nervous-system development, winter-lagged SNPs enriched for axonogenesis and cell projection morphogenesis, and summer-lagged SNPs for development of the fly’s excretory and osmoregulatory organs (Fig. 4D). Finally, genes containing signal for every combination of trigger season and response timing were again significantly enriched specifically for developmental and neural terms (Fig. 5E). The combined signal makes neural and developmental function the clearest, most consistently recovered pattern driving evolutionary change.

### Intersections of temporal modes highlight pleiotropic loci for stress resistance

Genes with both concurrent and lagged GEA signal are increasingly likely to have real phenotypic effects as pleiotropy increases. A genome-wide association study (GWAS) in inbred lines from our focal Pennsylvania population independently tested whether seasonal GEA SNPs have functional phenotypic effects on stress tolerance, specifically chill recovery, heat knockdown, and starvation resistance (fig. S6). We quantified pleiotropy as the number of traits for which a gene harbored at least one significant SNP (p<0.001); a score of one is not truly pleiotropic, while three means a gene affects all three traits tested. Both concurrent and lagged representation increased with the number of GWAS traits for which a gene was significant, from 89% and 59–65% respectively among single-trait genes to 100% among the 32 genes significant for all traits (Fig. 6A, fig. S7). The relationship is bidirectional with genes carrying both concurrent and lagged signal significantly enriched for GWAS hits (OR=3.18), while single-mode genes were significantly depleted (OR=0.35–0.47; Fig. 6B). This shows that the intersection of both temporal modes, but not either alone, tags the genomic positions where phenotypic effects on stress resistance are most concentrated. The 32 triple-overlap genes converged on six conserved functional pathways: ecdysone signaling, cAMP signaling, serotonin and ion channels, courtship and social behavior, cytoskeleton and motor proteins, and development and Hox genes (Fig. 6C, fig. S8). None of these are canonical stress-response or metabolic pathways; instead, they represent the neural, developmental, and behavioral machinery through which organisms sense and respond to their environment.

**Fig. 6.**
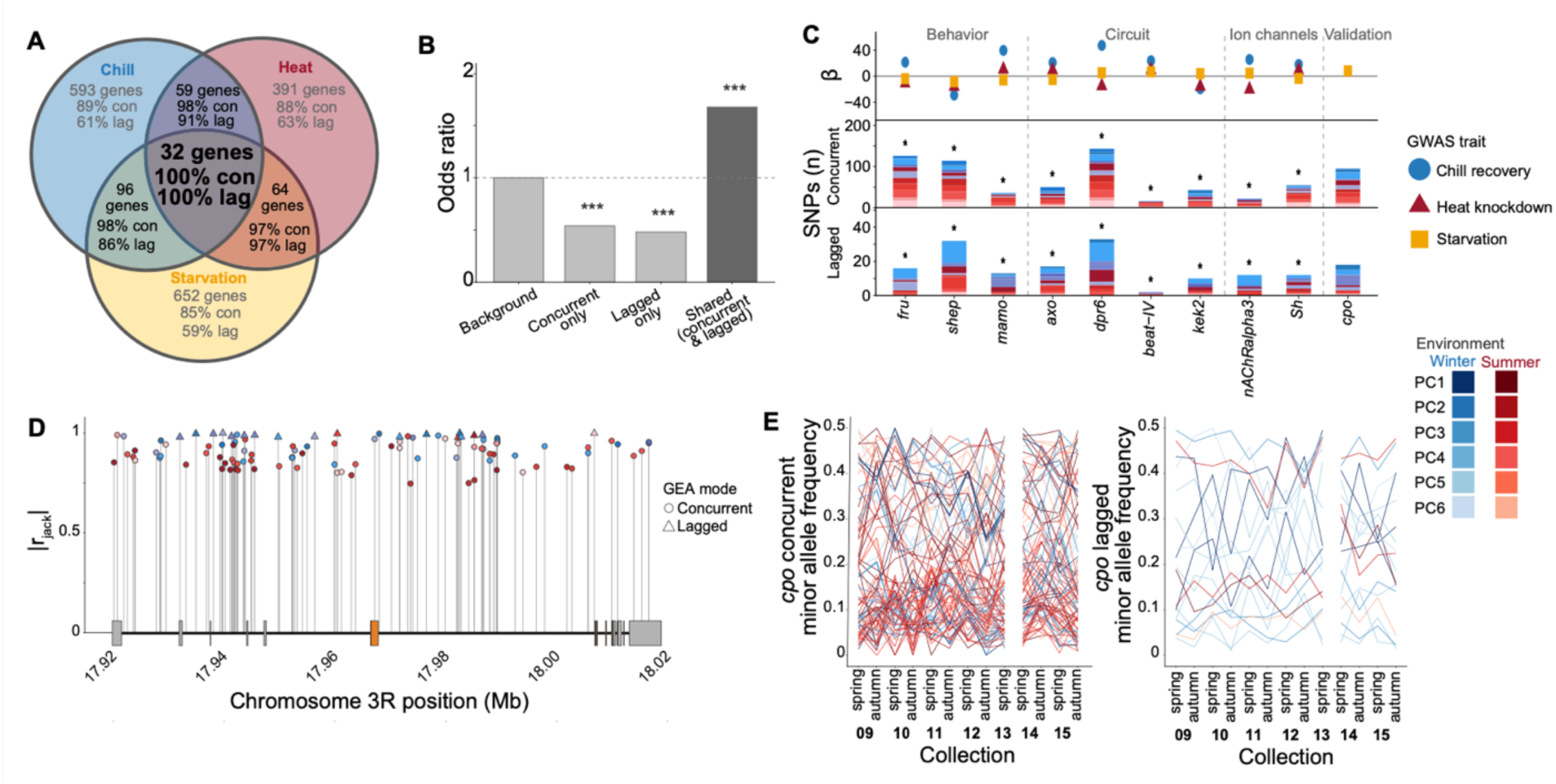
Pleiotropic genes linking stress-tolerance phenotypes to seasonal genotypic change converge across concurrent and lagged modes. **(A)** Genome-wide association (GWAS, P<0.001) using inbred lines from Pennsylvania population, for chill recovery, heat knockdown, and starvation resistance. 32 genes were pleiotropic across all three, with concurrent/lagged GEA overlap noted per trait and for the shared set. **(B)** Odds a gene is a GWAS hit for any trait, concurrent-only, lagged-only, and shared genes versus background. **(C)** Courtship/behavior genes among the 32 pleiotropic hits; points, GEA effect size (|r jack|), colored by GWAS trait; bars, GEA SNP counts, lagged (top) and concurrent (bottom), *couch potato (cpo),* not one of the 32 but included as independent validation given its established roles in seasonal adaptation, pleiotropy, and learning. **(D)** *cpo* gene structure with exons in orange and regulatory regions in gray. SNPs are plotted by location in the gene on x and significance on the y axis with mode indicated by shape and PC by color. **(E)** *cpo* minor allele frequency of GEA SNPs by selection mode.

As an independent check on our hit interpretation, we examined a gene already established as pleiotropic and relevant to seasonal evolution. *couch potato* (*cpo*) serves this purpose: it was not identified as one of the 32 genes above, and was a GWAS hit for only one of the three stress-tolerance traits tested here, yet it is independently well established in the seasonal adaptation literature for its role in dormancy, stress tolerance, and learning (*16–18*). *cpo* carries extensive concurrent and lagged GEA signal, with SNPs distributed across the full length of the gene rather than clustered in one region (Fig. 6D; gene structure for all 32 triple-overlap genes shown for comparison in fig. S9). These SNPs oscillate between spring and autumn each year, but different SNPs within the gene oscillate out of phase with one another (Fig. 6E), consistent with *cpo*’s pleiotropic effects arising from multiple, partially independent variants rather than one dominant haplotype.

## Discussion

A fundamental tension in evolutionary biology is the apparent unpredictability of adaptation in the wild: laboratory experiments often show repeatable trajectories, while field studies frequently report stochastic or undirected responses. We propose this tension reflects a central blind spot: the inability to distinguish a population’s immediate response to current selection from the genomic echo of a previous selective episode. Because most genomic studies rely on static snapshots, they cannot tell a response to the present from the residue of the past. Our findings address this directly: adaptation to recurring environmental heterogeneity proceeds through a temporally partitioned genomic framework, operating simultaneously across multiple timescales. This genomic framework is built from thousands of modest-effect loci that decouple the evolutionary response from immediate environmental triggers, with the circuitry linking environmental sensing to behavior emerging as a recurring node. The resulting hierarchy of responses, ranging from within-generation plasticity to multi-seasonal lagged genomic echoes, suggests a general strategy for how pleiotropic, polygenic traits evolve under structured environmental variability.

Severe selective episodes act as the organizing force at the center of this framework. In our system, winter serves as the dominant such episode, consistent with its role as the primary survival challenge of temperate seasonality. This dominance reflects environmental trigger and severity rather than timing: winter- and summer-triggered responses are organized around genuinely different environmental features, regardless of whether the outcome is immediate or carried into the following season. The result is a structured hierarchy of adaptation, with developmental plasticity managing within-season uncertainty while concurrent and lagged genomic modes supply the directional signals that connect populations across years (*19–21*).

Understanding how evolution unfolds in real time requires a commitment to long-term monitoring of wild populations. Experimental manipulation is an essential tool for validating specific hypotheses, yet it cannot capture the unscripted environmental variability that shapes natural populations over years. The lagged mode identified here illustrates this limitation: phenotypic carry-over is documented in other taxa (*7*), but uncovering a corresponding genomic signal required longitudinal tracking of a wild population. While the value of long-term studies has been recognized, the episodic nature of rare, severe climatic events often limits the analysis to showing that severity matters, without resolving how (*3*, *22–24*). The recurring structure of seasonal selection provides a higher-resolution lens; because winter severity varies continuously across independent years, we can determine not only if adaptation is predictable, but the degree to which it is. Our results reveal that predictability is not a fixed population trait, but a conditional state that scales continuously with selective severity, reflecting a tuned response to the intensity of the triggering environment rather than a binary threshold. This may help explain why previous long-term field studies have sometimes found evolution to be unpredictable (*25*); predictability itself may be an emergent property of how severe the selective episode is, not a fixed feature of the organism or system under study.

Several lines of evidence confirm that these patterns reflect genuine selection on standing genetic variation, rather than migration or demographic history. Concurrent and lagged SNPs show low haplotype linkage even where physically proximate, and parallel seasonal allele-frequency change across geographically distant populations indicates local selection rather than gene flow (*10*, *12*). That this signal is detectable at all is notable given the inherent limitations of pool-seq, which does not preserve haplotype structure and, combined with soft-sweep dynamics (*26*) and ongoing recombination in outbred populations (*27*), should weaken and diffuse adaptive signal. A signal spanning six environmental axes, two temporal modes, and thousands of independent SNPs despite these constraints speaks to the strength and repeatability of seasonal selection in this population.

The discovery of a lagged genomic mode, in which one season’s environment structures the allele frequencies of the next, raises the question of what mechanism sustains it. While the precise mechanism remains open, our data suggest that harsher environments impose stronger selection, which more strongly shapes the genetic composition of subsequent generations. This is supported by our phenotypic results, where winter severity determines the predictability and direction of subsequent summer tracking. Other mechanisms could also contribute. Long-distance epistasis may occur, in which alleles selected in one season alter the fitness landscape at unlinked SNPs in the next (*28*, *29*). Transgenerational plasticity is also possible, in which parental state directly shapes offspring phenotype (*30*, *31*). Life-stage carry-over more generally could contribute too, such as the lasting signals left on development by photoperiodic cues during diapause (*32*, *33*). Distinguishing among these mechanisms will require experimental manipulation, but the existence of a lagged mode suggests that selection is not simply a series of independent events, but can also unfold as a continuous trajectory.

This temporally partitioned framework provides a compelling explanation for the vast amount of adaptive genetic variation observed in natural populations. Single-axis models of selection often struggle to explain the high number of SNPs under selection without invoking implausibly strong selective pressures (*34*, *35*). However, our results show that independent environmental axes each drive distinct genomic responses with minimal locus sharing, effectively multiplying the number of genomic targets available to selection. When these multivariate axes are further multiplied by two independent temporal modes (concurrent and lagged) acting on largely non-overlapping SNPs, the result is a vast, layered genomic substrate for adaptation. This suggests that the search for a few major-effect alleles may be an unrepresentative approach to understanding evolutionary change (*36*). Instead, our work highlights a pervasive, polygenic signal where common, modest-effect polymorphisms are maintained by fluctuating selection to provide a flexible, tunable response to environmental heterogeneity (*37*).

Pleiotropy and epistasis together help explain how a limited number of loci can drive rapid, widespread adaptation. This is consistent with growing recognition that pleiotropy may be far more widespread among genetic variants than classical models assumed, though its true scope remains actively debated (*38*). Genes carrying both concurrent and lagged signal were significantly enriched for GWAS hits, directly tying this genomic overlap to real phenotypic effects: a single gene can influence multiple traits at once, consistent with experimental evidence that selection on behavior can act simultaneously with selection on other traits, such as body size and shape (*39*). Because concurrent and lagged variants often converge on the same genes, these variants may also interact epistatically, with combined effects exceeding what either produces alone. *couch potato* illustrates this architecture directly: it carries established roles in multiple life-history traits (*16*, *17*), alongside non-additive epistatic interactions among its own intronic SNPs that facilitate environmental adaptation (*17*).

Finally, we find that these temporal modes converge on a specific functional target: the neural and developmental circuitry by which organisms sense and respond to their environment. The genes most strongly linked to pleiotropic phenotypic effects converge on this same circuitry, suggesting it may be a recurring target of multi-temporal selection. Behavior, in particular, may act as both a target of selection and a buffer against environmental extremes (*40–43*). *fruitless*, the master regulator of sexually dimorphic neural circuitry and courtship behavior, satisfies our own screening criteria and illustrates this directly: a natural polymorphism in *fruitless* carries measurable fitness consequences in wild populations (*44*), consistent with the modulation of reproductive and social behavior itself being an ongoing target of selection. *couch potato*, despite falling outside our own screen’s criteria, provides independent support for the same pattern: it is already established in the literature for its role in dormancy, stress tolerance, and learning (*16–18*), strengthening the case that this convergence reflects a genuine biological signal rather than an artifact of our method. Reproductive preferences shaped by current conditions could also directly contribute to the lagged signal, by favoring genotypes suited to a passing context and thereby carrying that signal into the following season’s cohort.

Together, these results demonstrate that rapid adaptation does not proceed through a single dominant pathway. Instead, populations utilize a layered genomic architecture to integrate environmental information across time, ensuring survival in the face of an ever-shifting selective landscape.

## Methods

We combined longitudinal common-garden phenotypic data, pool-seq population genomics, and meteorological records from a single *Drosophila melanogaster* population in Pennsylvania sampled across seven years (2009–2015) (*10*, *12*). Four stress-tolerance and life-history traits were measured after four generations in a common laboratory environment to remove environmental effects, enabling inference of the genetic basis of seasonal phenotypic change. Seasonal allele frequency changes were correlated with six environmental PCs derived from meteorological data (NOAA) that collectively explain 99.4% of environmental variance using a jackknife-resampled Pearson correlation, yielding 65,403 unique concurrent GEA SNPs after LD pruning (r^2^>0.5, 50bp windows) and requiring each locus to exceed its own PC x season combination’s empirical permutation threshold. Lagged GEA correlated winter PC scores with the following summer’s allele frequency changes, and summer PC scores with the following winter’s allele frequency changes, using identical criteria, yielding 16,391 SNPs; spatial co-localization analysis showed these were modestly enriched near a concurrent locus at short range (≤25bp) but significantly depleted at longer range (≥50bp), indicating that lagged and concurrent SNPs are largely, though not entirely, genomically independent. Genome-wide association analysis for three stress-resistance traits in 123 inbred lines derived from the same population (*45*) identified candidate genes using linear models with season as covariate (p<0.001). SNPs within cosmopolitan inversion regions were flagged for separate enrichment analyses (*46*). Functional enrichment used goseq with gene-length correction (*47*). Broad functional categories (neural/behavior, development, life history, metabolism/stress, signaling, other) were assigned manually to provide a consistent classification scheme across all figures. To test whether these largely independent loci nonetheless converged on shared genes, gene-level overlap among the four seasonal categories was assessed against a permutation-based null and followed by functional enrichment analysis of the resulting convergent gene set. Full methods are provided in the Supplementary Information.

## Acknowledgments

We thank the owners and staff of Linvilla Orchards (Media, PA), George Hill Orchard (Lancaster, MA), and Carter Mountain Orchard (Charlottesville, VA) for permission to collect *Drosophila* on their properties. Meteorological data were provided by the NOAA National Centers for Environmental Information Global Historical Climatology Network Daily database. We thank Mark C. Bitter, Mark A. McPeek, Mary Lou Guerinot, and Carey D. Nadell for useful feedback on the manuscript. The authors used Claude (Anthropic) between May and October 2026 to assist with troubleshooting and streamlining analysis code originally written by the authors, and with editing and refining the language of the manuscript text. All analytical decisions, interpretation of results, and scientific conclusions are the authors’ own.

## Author contributions

E.L.B. and P.S. conceptualized the longitudinal seasonal study; E.L.B. conceptualized the genomic and environmental synthesis. E.L.B. and P.S. collected field samples; E.L.B. performed the experiments and all formal analyses and created all figures; E.L.B. wrote the manuscript; E.L.B. and P.S. reviewed and edited it. Funding was acquired by P.S., with additional support from E.L.B.

## Data and materials availability

Raw pool-seq sequencing reads underlying the concurrent and lagged genome-environment association analyses were previously published and are available from the NCBI Sequence Read Archive under BioProject accession PRJNA308584 (*10*, *12*), with processed allele frequency data and sample metadata available via the Dryad Digital Repository (https://doi.org/10.5061/dryad.4r7b826). Genome sequencing data for the 123 inbred lines used in the genome-wide association analysis are available from the NCBI Sequence Read Archive under BioProject accession PRJNA383555. The isofemale lines used for phenotypic assays are no longer available; the inbred lines used for genome-wide association analysis are maintained in the Schmidt and Behrman laboratories and are available upon reasonable request. The *D. melanogaster* reference genome assembly (BDGP6.32, Ensembl release 109) used for gene annotation is publicly available from Ensembl. Phenotype data, environmental data, and all analysis code generated for this study are provided as a supplementary file with this submission and will be made publicly available via Dryad and GitHub upon acceptance.

## Supplementary Materials

Materials and Methods

Table S1

Figs. S1 to S9

## References

1. N. Barghi, J. Hermisson, C. Schlötterer, Polygenic adaptation: a unifying framework to understand positive selection. Nat Rev Genet 21, 769–781 (2020).

2. P. de Villemereuil, A. Charmantier, D. Arlt, P. Bize, P. Brekke, L. Brouwer, A. Cockburn, S. D. Côté, F. S. Dobson, S. R. Evans, M. Festa-Bianchet, M. Gamelon, S. Hamel, J. Hegelbach, K. Jerstad, B. Kempenaers, L. E. B. Kruuk, J. Kumpula, T. Kvalnes, A. G. McAdam, S. E. McFarlane, M. B. Morrissey, T. Pärt, J. M. Pemberton, A. Qvarnström, O. W. Røstad, J. Schroeder, J. C. Senar, B. C. Sheldon, M. van de Pol, M. E. Visser, N. T. Wheelwright, J. Tufto, L.-M. Chevin, Fluctuating optimum and temporally variable selection on breeding date in birds and mammals. Proceedings of the National Academy of Sciences 117, 31969–31978 (2020).

3. P. R. Grant, B. R. Grant, Unpredictable Evolution in a 30-Year Study of Darwin’s Finches. Science 296, 707–711 (2002).

4. Bumpus, “The elimination of the unfit as illustrated by the introduced sparrow, Passer domesticus” in Biological Lectures Delivered at the Marine Biological Laboratory of Woods Hole in the Summer Session of 1897 (Boston: Ginn & Company, 1898), pp. 209–226.

5. T. Clutton-Brock, B. C. Sheldon, Individuals and populations: the role of long-term, individual-based studies of animals in ecology and evolutionary biology. Trends in Ecology & Evolution 25, 562–573 (2010).

6. C. Rellstab, F. Gugerli, A. J. Eckert, A. M. Hancock, R. Holderegger, A practical guide to environmental association analysis in landscape genomics. Molecular Ecology 24, 4348–4370 (2015).

7. X. A. Harrison, J. D. Blount, R. Inger, D. R. Norris, S. Bearhop, Carry-over effects as drivers of fitness differences in animals: Carry-over effects in animal populations. Journal of Animal Ecology 80, 4–18 (2011).

8. D. S. Saunders, V. C. Henrich, L. I. Gilbert, Induction of diapause in Drosophila melanogaster: photoperiodic regulation and the impact of arrhythmic clock mutations on time measurement. Proc. Natl. Acad. Sci. U.S.A. 86, 3748–3752 (1989).

9. E. L. Behrman, S. S. Watson, K. R. O’Brien, M. S. Heschel, P. S. Schmidt, Seasonal variation in life history traits in two *Drosophila* species. J. Evol. Biol. 28, 1691–1704 (2015).

10. A. O. Bergland, E. L. Behrman, K. R. O’Brien, P. S. Schmidt, D. A. Petrov, Genomic Evidence of Rapid and Stable Adaptive Oscillations over Seasonal Time Scales in Drosophila. PLoS Genet 10, e1004775 (2014).

11. M. Kapun, J. C. B. Nunez, M. Bogaerts-Márquez, J. Murga-Moreno, M. Paris, J. Outten, M. Coronado-Zamora, C. Tern, O. Rota-Stabelli, M. P. G. Guerreiro, S. Casillas, D. J. Orengo, E. Puerma, M. Kankare, L. Ometto, V. Loeschcke, B. S. Onder, J. K. Abbott, S. W. Schaeffer, S. Rajpurohit, E. L. Behrman, M. F. Schou, T. J. S. Merritt, B. P. Lazzaro, A. Glaser-Schmitt, E. Argyridou, F. Staubach, Y. Wang, E. Tauber, S. V. Serga, D. K. Fabian, K. A. Dyer, C. W. Wheat, J. Parsch, S. Grath, M. S. Veselinovic, M. Stamenkovic-Radak, M. Jelic, A. J. Buendía-Ruíz, M. J. Gómez-Julián, M. L. Espinosa-Jimenez, F. D. Gallardo-Jiménez, A. Patenkovic, K. Eric, M. Tanaskovic, A. Ullastres, L. Guio, M. Merenciano, S. Guirao-Rico, V. Horváth, D. J. Obbard, E. Pasyukova, V. E. Alatortsev, C. P. Vieira, J. Vieira, J. R. Torres, I. Kozeretska, O. M. Maistrenko, C. Montchamp-Moreau, D. V. Mukha, H. E. Machado, K. Lamb, T. Paulo, L. Yusuf, A. Barbadilla, D. Petrov, P. Schmidt, J. Gonzalez, T. Flatt, A. O. Bergland, Drosophila Evolution over Space and Time (DEST): A New Population Genomics Resource. Molecular Biology and Evolution 38, 5782–5805 (2021).

12. H. E. Machado, A. O. Bergland, R. Taylor, S. Tilk, E. Behrman, K. Dyer, D. K. Fabian, T. Flatt, J. González, T. L. Karasov, B. Kim, I. Kozeretska, B. P. Lazzaro, T. J. Merritt, J. E. Pool, K. O’Brien, S. Rajpurohit, P. R. Roy, S. W. Schaeffer, S. Serga, P. Schmidt, D. A. Petrov, Broad geographic sampling reveals the shared basis and environmental correlates of seasonal adaptation in Drosophila. eLife 10, e67577 (2021).

13. S. M. Rudman, S. I. Greenblum, S. Rajpurohit, N. J. Betancourt, J. Hanna, S. Tilk, T. Yokoyama, D. A. Petrov, P. Schmidt, Direct observation of adaptive tracking on ecological time scales in Drosophila. Science 375, eabj7484 (2022).

14. M. C. Bitter, S. Berardi, H. Oken, A. Huynh, E. Lappo, P. Schmidt, D. A. Petrov, Continuously fluctuating selection reveals fine granularity of adaptation. Nature 634, 389–396 (2024).

15. N. Barghi, R. Tobler, V. Nolte, A. M. Jakšić, F. Mallard, K. A. Otte, M. Dolezal, T. Taus, R. Kofler, C. Schlötterer, Genetic redundancy fuels polygenic adaptation in Drosophila. PLoS Biol 17, e3000128 (2019).

16. P. S. Schmidt, C.-T. Zhu, J. Das, M. Batavia, L. Yang, W. F. Eanes, An amino acid polymorphism in the *couch potato* gene forms the basis for climatic adaptation in *Drosophila melanogaster*. Proc. Natl. Acad. Sci. U.S.A. 105, 16207–16211 (2008).

17. E. L. Behrman, A. O. Bergland, D. Petrov, P. L. Schmidt, Intragenic epistasis in couch potato shapes life history traits and climatic adaptation in Drosophila melanogaster. bioRxiv [Preprint] (2026). 10.64898/2026.10.09.758124.

18. E. L. Behrman, T. J. Kawecki, P. Schmidt, Rapid evolution of learning and reproduction in natural populations of Drosophila melanogaster. bioRxiv [Preprint] (2026). 10.1101/288696.

19. N. G. Hairston, S. P. Ellner, M. A. Geber, T. Yoshida, J. A. Fox, Rapid evolution and the convergence of ecological and evolutionary time. Ecology Letters 8, 1114–1127 (2005).

20. F. Pelletier, D. Garant, A. P. Hendry, Eco-evolutionary dynamics. Philos Trans R Soc Lond B Biol Sci 364, 1483–1489 (2009).

21. G. Bell, Fluctuating selection: the perpetual renewal of adaptation in variable environments. Philos Trans R Soc Lond B Biol Sci 365, 87–97 (2010).

22. A. Charmantier, R. H. McCleery, L. R. Cole, C. Perrins, L. E. B. Kruuk, B. C. Sheldon, Adaptive Phenotypic Plasticity in Response to Climate Change in a Wild Bird Population. Science 320, 800–803 (2008).

23. A. Ozgul, S. Tuljapurkar, T. G. Benton, J. M. Pemberton, T. H. Clutton-Brock, T. Coulson, The Dynamics of Phenotypic Change and the Shrinking Sheep of St. Kilda. Science 325, 464–467 (2009).

24. D. Reznick, J. A. Endler, The Impact of Predation on Life History Evolution in Trinidadian Guppies (Poecilia Reticulata). Evolution 36, 160–177 (1982).

25. Z. D. Blount, R. E. Lenski, J. B. Losos, Contingency and determinism in evolution: Replaying life’s tape. Science 362, eaam5979 (2018).

26. J. Hermisson, P. S. Pennings, Soft Sweeps. Genetics 169, 2335–2352 (2005).

27. R. Barrett, D. Schluter, Adaptation from standing genetic variation. Trends in Ecology & Evolution 23, 38–44 (2008).

28. A. M. Taverner, L. Yang, Z. J. Barile, B. Lin, J. Peng, A. P. Pinharanda, A. S. Rao, B. P. Roland, A. D. Talsma, D. Wei, G. Petschenka, M. J. Palladino, P. Andolfatto, Adaptive substitutions underlying cardiac glycoside insensitivity in insects exhibit epistasis in vivo. eLife 8, e48224 (2019).

29. C. Bank, Epistasis and Adaptation on Fitness Landscapes. Annu. Rev. Ecol. Evol. Syst. 53, 457–479 (2022).

30. T. Uller, Developmental plasticity and the evolution of parental effects. Trends in Ecology & Evolution 23, 432–438 (2008).

31. R. Bonduriansky, T. Day, Nongenetic Inheritance and Its Evolutionary Implications. Annu. Rev. Ecol. Evol. Syst. 40, 103–125 (2009).

32. Z. Tian, K. Wang, S. Guo, J.-X. Li, K. King-Jones, F. Zhu, W. Liu, X.-P. Wang, The PBAP chromatin remodeling complex mediates summer diapause via H3K4me3-driven juvenile hormone regulation in *Colaphellus bowringi*. Proc. Natl. Acad. Sci. U.S.A. 122, e2422328122 (2025).

33. L. Salis, E. Van Den Hoorn, D. G. M. Beersma, R. A. Hut, M. E. Visser, Photoperiodic cues regulate phenological carry-over effects in an herbivorous insect. Functional Ecology 32, 171–180 (2018).

34. V. Buffalo, G. Coop, Estimating the genome-wide contribution of selection to temporal allele frequency change. Proc. Natl. Acad. Sci. U.S.A. 117, 20672–20680 (2020).

35. O. L. Johnson, R. Tobler, J. M. Schmidt, C. D. Huber, Fluctuating selection and the determinants of genetic variation. Trends in Genetics 39, 491–504 (2023).

36. M. V. Rockman, The QTN Program and the Alleles That Matter for Evolution: All That’s Gold Does Not Glitter. Evolution 66, 1–17 (2012).

37. M. C. Bitter, P. Schmidt, D. Petrov, Common, consequential, and actively maintained: a neo-balance view of genetic variation. EcoEvoRxiv [Preprint] (2026). https://ecoevorxiv.org/repository/view/15208/.

38. A. B. Paaby, M. V. Rockman, The many faces of pleiotropy. Trends Genet 29, 66–73 (2013).

39. O. Lapiedra, T. W. Schoener, M. Leal, J. B. Losos, J. J. Kolbe, Predator-driven natural selection on risk-taking behavior in anole lizards. Science 360, 1017–1020 (2018).

40. E. Mayr, “The emergence of evolutionary novelties” in Evolution after Darwin. Vol. 1. The Evolution of Life, S. Tax, Ed. (Chicago, 1960), pp. 349–380.

41. W. T. Wcislo, Behavioral Environments and Evolutionary Change. Annual Review of Ecology and Systematics 20, 137–169 (1989).

42. M. J. West-Eberhard, Developmental Plasticity and Evolution (Oxford University Press, Oxford, New York, 2003).

43. R. A. Duckworth, The role of behavior in evolution: a search for mechanism. Evol Ecol 23, 513–531 (2009).

44. M. D. Jardine, F. Ruzicka, C. Diffley, K. Fowler, M. Reuter, A non-coding indel polymorphism in the fruitless gene of Drosophila melanogaster exhibits antagonistically pleiotropic fitness effects. Proc Biol Sci 288, 20202958.

45. E. L. Behrman, V. M. Howick, M. Kapun, F. Staubach, A. O. Bergland, D. A. Petrov, B. P. Lazzaro, P. S. Schmidt, Rapid seasonal evolution in innate immunity of wild Drosophila melanogaster. Proceedings of the Royal Society B: Biological Sciences 285, 20172599 (2018).

46. M. Kapun, D. K. Fabian, J. Goudet, T. Flatt, Genomic Evidence for Adaptive Inversion Clines in *Drosophila melanogaster*. Mol Biol Evol 33, 1317–1336 (2016).

47. M. D. Young, M. J. Wakefield, G. K. Smyth, A. Oshlack, Gene ontology analysis for RNA-seq: accounting for selection bias. Genome Biol 11, R14 (2010).

48. V. Trotta, F. C. Calboli, M. Ziosi, D. Guerra, M. C. Pezzoli, J. R. David, S. Cavicchi, Thermal plasticity in Drosophila melanogaster: a comparison of geographic populations. BMC Evol Biol 6, 67 (2006).

49. C. D. Schlichting, The Evolution of Phenotypic Plasticity in Plants. Annual Review of Ecology and Systematics 17, 667–93 (1986).

