## supplements for "Evolution Tracks Multiple Timescales at Once"

### Materials and Methods

#### Study populations and field sampling

Wild *Drosophila melanogaster* were collected using aspiration from June 2009 through November 2015 at Linvilla Orchards in Media, PA (39.884°N, 75.411°W). Isofemale lines established from collections beginning June 2011 were retained for phenotypic assays; all collections from 2009 onward were used for pool-seq genomic analysis (see below). Additional collections were made in July and October of 2012 and 2014 at locations approximately equidistant north and south: George Hill Orchard in Lancaster, MA (42.500°N, 71.564°W) and Carter Mountain Orchard in Charlottesville, VA (37.992°N, 78.472°W). Collection targets aimed for approximately 100 isofemale lines per collection point, constrained by the number of gravid wild-caught females successfully establishing lines; between 18 and 106 isofemale lines were established per collection (mode of 40, table S1). Flies were reared on cornmeal-molasses food under standard laboratory conditions (25°C, 12L:12D).

Inbred lines were generated from spring and autumn isofemale lines collected in Pennsylvania in 2012. The lines were inbred by full-sib mating for 20 generations and subsequently sequenced (NCBI SRA BioProject PRJNA383555) (*43*). No genetic modifications were introduced in any isofemale or inbred lines; all lines are wild-derived from field-caught *Drosophila melanogaster*.

#### Common garden phenotype assays

Four life-history and stress-tolerance traits were measured after four generations in a common laboratory environment (25°C, 12L:12D): developmental time (egg-to-adult), chill coma recovery time (minutes to resume locomotion after cold paralysis), heat knockdown resistance (minutes to knockdown), and starvation resistance (hours to death on water only). Rearing for four generations in a common environment substantially removes transgenerational environmental effects, so phenotypic differences among collections are inferred to reflect primarily genetic rather than environmentally driven differences in traits.

Developmental time was quantified as time from oviposition to adult emergence; a single gravid female was placed into a vial for 24 hours to standardize egg density, with ten replicate vials per isofemale line. Starvation resistance was assessed for 12 individuals per sex per line measured in groups. Flies were briefly anesthetized with $CO_2$, allowed to recover for 24 hours on food, then transferred to vials with water-only access where survival was scored at three standardized timepoints per day. DAM2 activity data were additionally filtered to exclude false-positive signals: chill coma recovery times under 2 minutes were excluded, as these likely reflect spurious infrared beam breaks rather than genuine locomotor recovery. Chill coma recovery was capped at a maximum scored time of 2 hours and heat knockdown at 3 hours. No upper or lower time caps were applied in the thermal plasticity assay.

Thermal tolerance assays used DAM2 activity monitors (TriKinetics, Waltham, MA) to record locomotor activity of 3–5 day old males and females separately at 10-second intervals (*9*). Chill coma recovery was quantified as time to resume locomotor activity at 25°C following 2 hours of cold paralysis at 4°C. Heat knockdown was measured as time to cessation of locomotor activity during a temperature ramp from 25°C increasing at 1°C per minute to 37°C, held constant at that temperature thereafter. Flies were randomly assigned to DAM2 monitor channels within each assay run. Which isofemale lines were assayed together in a given run was determined by the availability of sufficient age-matched flies at the same time, not by randomization or any trait-related criterion; line numbers reflect only the arbitrary order lines were established and carry no experimental meaning. DAM2 monitor assays (chill coma recovery, heat knockdown) were collected automatically by the instrument, blind to line identity. Starvation resistance and developmental time were scored manually and were not blinded; however, line identity could not systematically bias scoring for the reason above.

### Generation model

The number of generations elapsed between seasonal collections was estimated using a degree-day accumulation model. Daily mean temperature was calculated as the mean of daily maximum and minimum temperatures. The active growing season was defined as days when both the 3-day rolling mean temperature exceeded 12°C and daylength exceeded 11 hours, thresholds consistent with published data on dormancy induction and termination in *D. melanogaster* (*8*). Degree days were accumulated during active periods using a base temperature of 12°C and a ceiling of 27°C, above which development rate was held constant rather than continuing to increase with temperature — an estimate that likely undercounts true generation number, since warmer days are capped rather than credited with the faster development they would actually produce. Daylength was calculated from latitude and day of year using the geosphere package in R. One generation was defined as 112 accumulated degree days, based on published temperature-development relationships for *D. melanogaster* (*48*). Cumulative and within-year generation counts were calculated separately for each population and year.

### Environment data and principal component analysis

Daily meteorological data were obtained from the Global Historical Climatology Network (NOAA) at weather stations nearest each collection site: Philadelphia International Airport (PA), Norwood Memorial Airport (MA), and Charlottesville-Albemarle Airport (VA). Variables included daily maximum and minimum temperature (°C), precipitation (mm), snowfall (mm), average wind speed (m/s), and wind direction (degrees). Wind direction was decomposed into east-west (sine) and north-south (cosine) components prior to analysis to account for its circular nature, yielding seven variables in total. Photoperiod was not included in the PCA despite its well-established role in *Drosophila* seasonal biology: because daylength at a given latitude is determined by date, mean seasonal photoperiod has near-zero year-to-year variance within a season and would contribute nothing to the between-year environmental variation that drives genome-environment association. Daylength was instead used to define the boundaries of the active growing season in the generation model (see above).

Environmental PCA was performed on mean seasonal values of these seven variables for the Pennsylvania site, which provided the complete longitudinal genomic time series. PCA was conducted using prcomp in R with centering and unit-variance scaling. Data were partitioned into winter (November–May) and summer (June–October) seasons; the winter-year was assigned as the calendar year of the following spring, so that the winter spanning 2011–12 is labeled 2012. Six principal components were retained, collectively explaining 99.4% of total environmental variance. PC1 (33.0%) captured temperature variation (dominant loadings: Tmax, Tmin); PC2 (17.8%) captured overall moisture input, with positive loadings for precipitation, snowfall, and the north-south wind component; PC3 (15.3%) captured high-speed wind conditions, with strong positive loadings for wind speed and strong negative loadings for the north-south wind component; PC4 (13.6%) captured wind and dryness, with strong positive loadings for the east-west wind component and strong negative loadings for snowfall, reflecting cold dry air masses with suppressed precipitation; PC5 (11.2%) captured the rain-versus-snow precipitation axis, with positive loadings for liquid precipitation and negative loadings for snowfall, reflecting variation in temperatures above versus below freezing; and PC6 (8.4%) captured high-speed cross-directional wind conditions, with strong positive loadings for wind speed and the north-south wind component, and negative loadings for the east-west wind component. Biplots for all pairwise combinations of the six retained PCs are shown in fig. S2.

Winter severity was classified using environmental PCA scores. Mild winters were identified as those whose seasonal PC scores deviated from other winters toward the summer cluster in PC1–PC2 space, reflecting warmer minimum temperatures, reduced snowfall, and lower wind indices. This classification identified 2012 and 2013 as mild winters and 2009–2011 and 2014–2015 as harsh, and was confirmed by inspection of raw seasonal meteorological variables (fig. S1 and fig. S2).

Winter and summer environmental PC scores showed no significant same-PC or cross-PC correlation across the seven years sampled (all $p_{adj} > 0.05$, BH-corrected across 36 comparisons), indicating that cross-season concurrent GEA associations reflect independent seasonal signals rather than shared year-to-year environmental covariation. The same approach applied within each season, also across all seven sampled years, similarly found no significant covariation among the six PCs (all $p_{adj} > 0.05$, BH-corrected across 30 comparisons), indicating that the six environmental PCs behave as largely independent axes both within and across seasons.

**Statistical analyses of phenotype data**

Trait values were standardized to z-scores within trait and year prior to all phenotypic analyses. All tests used Benjamini-Hochberg (BH) correction for multiple comparisons across traits.

***Temporal phenotypic variation***

Phenotypic differences between adjacent seasonal collections were tested using Wilcoxon rank-sum tests on line means, with BH correction applied across all consecutive collection pairs and traits.

***Adaptive tracking***

To test whether traits tracked cumulative generations across the season, linear mixed models of the form trait ~ generations + (1|Year) were fit separately for collections following harsh and mild winters using the lme4 and lmerTest packages in R. Significance was assessed using the Satterthwaite approximation for degrees of freedom. Adaptive tracking analyses were restricted to the Pennsylvania population, which provided the complete longitudinal time series.

***Bet-hedging***

Within-collection phenotypic variance was quantified as a bet-hedging index using the coefficient of variation ($\sigma/\mu$) of individual replicate measurements for each isofemale line within a single collection (minimum 5 replicates per line per collection); each isofemale line exists only within one collection, so this CV is computed within a collection, never across years. Per-line CVs were then averaged to the collection level (one value per trait × collection). Two comparisons were made: (1) collections were classified as harsh or mild winter severity, and mean line CV was compared between severity classes by t-test; (2) collections were classified as spring (May–June) or fall (October–November), and mean line CV was compared between seasons by t-test. Both comparisons used BH correction across traits.

***Thermal plasticity***

Plasticity was assessed using spring and fall collections from 2012 (a mild year). Ten females from each of approximately 40 isofemale lines laid eggs for 24 hours at each of three larval rearing temperatures (18°C, 25°C, 29°C) under 12L:12D. Adults aged 3–5 days were then assayed for chill recovery, heat knockdown, and starvation resistance across the same three adult acclimation temperatures, yielding a fully factorial design with nine temperature combinations (two conditions missing for heat knockdown due to experimental loss). Plasticity indices were calculated as the coefficient of variation across temperature means per isofemale line, computed separately for larval (developmental acclimation) and adult (short-term acclimation) components, and combined as their mean (total plasticity index) (*49*). Seasonal differences between spring and autumn in plasticity index were tested by t-test with BH correction.

### Pool-seq and allele frequency data

Pooled whole-genome sequencing data, comprising one spring and one autumn pooled sample per year, were obtained from previously published datasets generated from Pennsylvania collections spanning 2009–2015 (*10*, *12*), with each sample comprising approximately 50–100 wild-caught males sequenced to an average read depth of 20–200x coverage. One autumn 2013 sample was excluded from all genomic analyses due to contamination with *D. simulans* (*12*), and one November 2011 sample was excluded from seasonal delta analyses as it represented an atypical frost collection rather than a standard spring or fall timepoint. This yielded 13 seasonal samples retained for analysis across five major chromosomal arms (2L, 2R, 3L, 3R, X).

Raw allele frequencies were extracted for 3,625,035 SNPs. Prior to analysis, SNPs were filtered to retain those with mean minor allele frequency ≥ 0.05 across all samples, no more than two samples with MAF < 0.02 (near-monomorphic), and no more than two samples with missing data. SNPs with functionally zero allele frequency change (delta ≈ 0, reflecting the precision of the reported frequency estimates) in more than two seasonal transitions were additionally excluded as uninformative. These filters retained 1,417,437 SNPs for downstream analysis. SNPs within known cosmopolitan inversion regions (In(2L)t, In(2R)NS, In(3L)P, In(3R)Mo, In(3R)Payne, In(3R)C) were identified using published breakpoint coordinates (*44*) and flagged for separate enrichment analyses.

#### *Concurrent genome-environment association*

Seasonal allele frequency change (delta) was calculated as the difference in allele frequency between consecutive spring and fall collections, yielding 5 winter deltas (autumn-to-spring transitions: W2010, W2011, W2012, W2013, W2015) and 6 summer deltas (spring-to-autumn transitions: S2009, S2010, S2011, S2012, S2014, S2015). Genome-environment associations were estimated as Pearson correlations between per-SNP seasonal deltas and each of the six environmental PC scores across corresponding years, computed separately for winter and summer seasons. The six PCs were derived from a single PCA fit to seasonal mean environmental data at the Pennsylvania site only, pooling winter and summer seasonal means together; the resulting PC loadings, which define what each axis represents environmentally, are therefore shared across both seasons, while each season-year's score on these shared axes is season-specific. Concretely, each SNP's seasonal allele-frequency change is a vector with one entry per seasonal transition (length 5 for winter, 6 for summer); this vector was correlated, per SNP and per PC, against the vector of that PC's season-specific score in the corresponding years — a SNP's winter deltas were correlated against winter-year PC scores, and its summer deltas against summer-year PC scores, using the same six PC definitions throughout. A significant correlation therefore indicates that a SNP's year-to-year seasonal trajectory tracks year-to-year variation in whichever environmental dimension that specific PC captures, within that season.

To account for the skewed sampling distribution of Pearson's r near ±1, correlations were transformed using Fisher's z-transformation prior to jackknife resampling. For each SNP and environmental PC, a leave-one-out jackknife estimated bias-corrected correlation coefficients and standard errors on this transformed scale. 95% confidence intervals were computed on the transformed scale and converted back to a correlation using the inverse transformation. A locus was considered significantly associated with a given environmental PC if its jackknife-derived 95% confidence interval excluded zero, the primary significance criterion, and its jackknife-estimated absolute correlation exceeded the empirical permutation threshold specific to that PC and season, a secondary magnitude criterion defined as the 95th percentile of a null distribution generated by shuffling PC scores relative to the observed allele-frequency trajectories, with 150 permutations per combination of PC and season.

To reduce redundancy from SNPs in high linkage disequilibrium, significant loci were pruned using genotype data from 123 lines derived from the same Pennsylvania population. SNPs were ranked by |r jack| and processed in descending order; where any two SNPs within 50bp had

$r^2$>0.5, the lower-ranked SNP was removed. Empirical analysis of the current set of significant loci confirmed that $r^2$ remains well below the 0.5 pruning threshold at all tested physical scales (median $r^2$=0.084 at <50bp, 0.146 at 50–100bp, 0.052 at 200–500bp; fig. S3), supporting the use of a 50bp pruning window. This retained 65,403 unique concurrent GEA loci across all PC × season combinations.

To test whether the winter–summer asymmetry in jackknife-corrected correlation strength reflected the difference in the number of seasonal transitions available for each season (five for winter, six for summer) rather than a genuine biological difference, we recomputed genome-wide jackknife |r| for summer after downsampling to five transitions, using all six possible five-of-six subsets (leave-one-transition-out) and averaging |r| per SNP across subsets. This comparison was performed genome-wide rather than restricted to already-significant loci, since loci pre-selected for extreme |r| under the original six-transition analysis would be expected to show reduced |r| under resampling for purely statistical reasons (regression to the mean), independent of any true seasonal difference.

#### *PC loading validation using raw environmental variables*

To confirm that GEA signal reflected genuine covariation with the underlying environmental axis rather than an artifact of the PCA reduction itself, jackknife correlations were recomputed between per-SNP allele-frequency deltas and the raw environmental variables contributing to each PC's dominant loadings (e.g., maximum and minimum temperature, temperature variance, and counts of extreme hot/cold days for PC1; wind speed and its variance and extreme-day counts, and the north-south wind component, for PC3), rather than the PC score itself. For each raw variable, this comparison was made separately for GEA-significant loci and a randomly sampled set of non-significant loci from the same PC × season combination, and separately for concurrent and lagged associations. As in the genome-wide winter–summer comparison described above, summer's raw-variable correlations were computed via leave-one-transition-out averaging to match winter's effective sample size of five transitions.

#### Lagged genome-environment association

Lagged GEA tests whether one season's environmental conditions predict allele-frequency change in the following season, rather than the same season's own environment (the concurrent analysis above); for example, a winter delta (spring minus preceding autumn) is here correlated against the preceding summer's PC scores, rather than that winter's own conditions. To confirm that lagged loci represent genuinely independent genomic variants rather than loci in linkage disequilibrium with concurrent GEA loci, we tested for spatial co-localization between the two sets. Of the two lagging directions, 8.7% of winter-lagged loci and 8.0% of summer-lagged loci were shared exactly with a concurrent locus. Among the remaining, non-exactly-shared loci, co-localization with a concurrent locus showed a distance-dependent pattern relative to chance expectation given the effective *Drosophila* genome size of ~40Mb, representing the mappable euchromatic portion of the genome after excluding heterochromatic and repetitive regions: lagged loci were significantly enriched for a nearby concurrent locus at short range (≤25bp; binomial test, all $p<0.002$), not significantly different from chance at 30–40bp, and significantly depleted at longer range (≥50bp; all $p<0.001$; fig. S4). To further test whether physically proximate concurrent and lagged loci might nonetheless represent the same causal variant captured by linkage disequilibrium, haplotype $r^2$ was computed between all concurrent–lagged SNP pairs within 500bp using genotype data from the Pennsylvania inbred lines. The median $r^2$ among pairs within 50bp was 0.267, below the LD pruning threshold of $r^2$=0.5. Together, these results indicate that while a small subset of lagged loci sit unusually close to a concurrent locus, the majority of lagged loci show no evidence of physical proximity to concurrent loci at the distances most relevant to linkage, and even physically proximate pairs are rarely in strong linkage.

### Seasonal routing network analysis

We constructed a routing network to characterize the architecture of cross-season genomic associations. Nodes represent environmental PC × season combinations and edges represent loci meeting the same significance criteria described above independently for each of the two connected PC × season combinations. For each pair of PC × season nodes, the number of shared loci was counted and the predominant direction of association was determined by comparing locus counts in each direction: pairs where more loci associated autumn→spring than spring→autumn (ratio ≥ 1.5) were classified as Winter→Summer directed edges; pairs with the reverse ratio (≤ 0.67) as Summer→Winter directed; and pairs with intermediate ratios as symmetrical. Each PC pair was represented by a single directed edge to avoid double-counting. Within-season edges connecting loci significant for multiple PCs within the same season were constructed separately. Edge width was proportional to the number of shared loci and node size was proportional to locus count, both scaled consistently using global min/max values across all panels.

Whether the observed proportion of loci associated with multiple environmental PCs within a season exceeded chance expectation was tested using an empirical permutation null: for each season, PC labels were randomly shuffled among that season's significant loci (2,000 iterations), preserving each locus's original number of PC assignments while reshuffling which specific PCs those assignments were drawn from. The proportion of loci associated with more than one PC was recalculated at each iteration, and observed multi-PC rates were compared against this permuted null distribution to obtain an empirical p-value. Winter and summer multi-PC rates were additionally compared to each other using a chi-square test with Yates' continuity correction.
Whether cross-season, cross-PC associations were genuinely asymmetric rather than reflecting sampling noise around an equal split was assessed by testing each asymmetric PC pair (edge ratio exceeding the 1.5-fold threshold in either direction) individually against a null of equal representation in both directions, using a two-sided binomial test with Benjamini-Hochberg correction applied across all tested pairs.

### Gene annotation and functional enrichment

Significant SNPs were mapped to genes by extending each annotated gene body 1 kb upstream in the *D. melanogaster* BDGP6.32 genome assembly (Ensembl release 109), loaded via the rtracklayer package. Overlapping SNPs were identified using the GenomicRanges package. Functional enrichment analysis was performed using the goseq package (*45*) with biological process (BP) ontology terms, correcting for gene length bias using the Wallenius approximation. Terms with BH-adjusted p ≤ 0.05 and a minimum of five focal genes per term were retained. The background gene set comprised all genes harboring at least one SNP in the post-filter genome-wide dataset. Functional terms were manually curated into six broad functional categories (neural/behavior, development, life history, metabolism/stress, signaling, other) with intergenic loci tracked separately, providing a consistent classification scheme across all figures. Separately from GO-term enrichment testing, individual loci were assigned to one of the six broad functional categories directly, for use in the functional-composition null checks described below (see Results). Each significant SNP was first mapped to its overlapping or nearest gene(s) as above. Each gene was then assigned a category by majority vote across its annotated GO terms, using the same fixed term-to-category lookup used to curate the six broad categories above; genes whose GO terms did not map to any category in this lookup were classified as "other," and loci with no overlapping gene were classified as "intergenic." Where a locus overlapped multiple genes with different assigned categories, the locus was assigned the category of the majority of its overlapping genes.

#### *Cross-category gene overlap*

Genes were mapped from each of the four seasonal categories (winter-concurrent, summer-concurrent, winter-lagged, summer-lagged) using the same gene-annotation procedure described above. The size of the four-way intersection, along with all pairwise and triple intersections, was computed directly from these gene sets. To test whether the four-way overlap exceeded chance expectation, a null distribution was generated by resampling genes directly, weighted by gene length, from the background gene universe (all genes appearing in any of the four categories, plus all genes with a known length in the reference annotation). For each of 200 permutations, four random gene sets were drawn matching each category's real observed gene count, and the size of their four-way intersection was recorded. Length-weighting corrects for the fact that longer genes have a higher probability of containing a significant SNP by chance alone, which would otherwise inflate apparent convergence among randomly drawn gene sets independent of any true biological signal.

**Inbred line genome-wide association study**

Genome-wide association analysis was performed using the 123 inbred lines derived from the 2012 Pennsylvania collections (*43*). Three stress-tolerance traits were analyzed: chill coma recovery time, heat knockdown resistance, and starvation resistance. Trait means were calculated per line by averaging sex-specific means, requiring a minimum of three individuals per sex per line. SNPs were retained with MAF $\geq 0.05$ and missingness $\leq 20\%$; missing genotypes were imputed to the per-SNP mean prior to association testing. Association was tested using a linear model of the form: phenotype ~ genotype + season, where season (spring or autumn collection) was included as a fixed covariate to account for systematic phenotypic differences between collection seasons. Significance was assessed using a $p < 0.001$ threshold to maximize power for identifying genes with consistent associations across all three traits simultaneously; the requirement for triple-overlap across independent traits provides an effective multiple-testing correction. These three traits are all components of climatic stress tolerance, so some degree of shared genetic architecture is plausible *a priori*. However, phenotypic correlations among the traits are weak, so the triple-overlap criterion is not simply recovering an expected correlated-trait signature, and the genes meeting it represent a conservative rather than a trivially expected result. Genes were considered GWAS hits if they harbored at least one SNP with $p < 0.001$ for all three traits simultaneously (triple-overlap criterion; 32 genes). Genes were classified into three mutually exclusive categories according to their GEA signal: concurrent-only, lagged-only, and shared (containing both concurrent and lagged loci). Enrichment for GWAS hits in each category was tested against the background of the other two categories combined (e.g., for the shared-gene comparison, the background was the union of concurrent-only and lagged-only genes), so that each of the three focal categories was tested against the genes in the remaining two categories combined.

**Pathway analysis**

Six biological pathways were identified based on the functional roles of the 32 triple-overlap genes (significant in all three GWAS traits and the concurrent GEA). A pathway was defined as a set of genes sharing an established, functionally related biological role, identified by reviewing the known function of each of the 32 genes individually and grouping those that shared a common biological theme. For each pathway, additional members were identified from published functional annotations in FlyBase and the primary literature. These pathway-expanded gene sets were then queried against the concurrent GEA, lagged GEA, and GWAS results to characterize the breadth of seasonal genomic associations within each pathway. For each pathway gene, concurrent GEA loci were counted separately for each PC × season combination and plotted as stacked bars colored by PC × season using the same color scheme as Fig. 3. Lagged GEA loci were counted separately by direction, summer-lagged (W→S), winter-lagged

(S→W), and PC. GWAS effect sizes (β) for the most significant SNP per gene per trait are shown above the GEA bars. Genes among the 32 triple-overlap set are indicated with asterisks.

***Statistical software***

All statistical analyses were performed in R (v4.3; R Core Team 2024), with Claude (Anthropic) assisting with debugging and reformatting analysis code; all statistical analyses, results, and interpretations were performed and verified by the authors. Key packages used included: lme4 (v2.0.1) and lmerTest for mixed models; goseq (v1.64.0) for functional enrichment corrected for gene length bias using the Wallenius approximation; GenomicRanges (v1.64.0) and VariantAnnotation (v1.58.0) for genomic data handling; ggrastr (v1.0.2) for rasterizing high-density point layers in genome-wide plots; ggforce, ggpubr, patchwork, and cowplot for figure construction and statistical annotation; ggplot2 (v4.0.3) for visualization; dplyr (v1.2.1) and purrr (v1.2.2) for data manipulation.

Extended Data Table 1. Collection dates of seasonal samples from Linvilla Orchard, Media, PA, George Hill Orchard, Lancaster, MA and Carter Mountain Orchard, Charlottesville, VA. Mean, standard error (SE) and number of isofemale lines sampled (n) are indicated for each collection.

| | | | | Development | | | Starvation | | | Chill | | | Heat | | |
|---|---|---|---|---|---|---|---|---|---|---|---|---|---|---|---|
| Year | Month | Date | Pop | mean | SE | n | mean | SE | n | mean | SE | n | mean | SE | n |
| 2011 | June | 1 | PA | 175.68 | 1.71 | 61 | 97.77 | 1.48 | 69 | 35.55 | 1.81 | 68 | 112.64 | 13.52 | 62 |
| 2011 | July | 31 | PA | 222.79 | 3.03 | 65 | 93.42 | 1.57 | 106 | 37.49 | 1.83 | 82 | 40.08 | 2.31 | 78 |
| 2011 | September | 26 | PA | 199.97 | 2.56 | 52 | 89.02 | 1.83 | 54 | 34.52 | 2.14 | 51 | 58.42 | 3.16 | 48 |
| 2011 | November | 9 | PA | 246.13 | 2.28 | 53 | 59.39 | 1.11 | 44 | 43.34 | 3.12 | 44 | 49.88 | 2.23 | 44 |
| 2012 | June | 1 | PA | 209.05 | 3.17 | 38 | 81.42 | 1.19 | 40 | 43.59 | 2.65 | 38 | 21.00 | 0.62 | 5 |
| 2012 | July | 19 | PA | 220.05 | 2.65 | 26 | 75.24 | 2.14 | 40 | 49.54 | 5.17 | 32 | 42.19 | 3.22 | 39 |
| 2012 | July | 18 | VA | 233.20 | 4.43 | 29 | 71.88 | 1.43 | 40 | 42.72 | 5.17 | 23 | 41.19 | 3.27 | 31 |
| 2012 | July | 31 | MA | 212.61 | 1.94 | 30 | 73.18 | 1.72 | 40 | 53.74 | 3.50 | 20 | 46.57 | 2.71 | 35 |
| 2012 | October | 10 | PA | 215.98 | 2.05 | 35 | 105.01 | 3.92 | 27 | 37.41 | 3.01 | 31 | 56.06 | 4.45 | 37 |
| 2012 | October | 12 | VA | 216.89 | 2.59 | 28 | 97.39 | 2.60 | 28 | 43.60 | 3.64 | 28 | 47.08 | 4.46 | 30 |
| 2012 | October | 14 | MA | 214.64 | 1.95 | 39 | 95.15 | 3.17 | 31 | 36.41 | 2.12 | 31 | 62.51 | 4.63 | 39 |
| 2012 | November | 7 | PA | 261.27 | 3.26 | 37 | 100.15 | 2.04 | 38 | 50.62 | 3.81 | 36 | 48.90 | 3.84 | 36 |
| 2013 | May | 13 | PA | 200.33 | 4.83 | 16 | 67.87 | 2.99 | 18 | - | - | - | 58.43 | 6.55 | 16 |
| 2013 | May | 30 | PA | 187.63 | 2.25 | 61 | 95.62 | 2.05 | 59 | 29.18 | 2.03 | 61 | 37.99 | 4.16 | 16 |
| 2013 | September | 6 | PA | 243.98 | 1.44 | 58 | 69.20 | 1.03 | 60 | 45.04 | 1.99 | 55 | 44.42 | 2.23 | 55 |
| 2013 | November | 8 | PA | 244.42 | 2.35 | 44 | 66.95 | 1.50 | 49 | 29.73 | 1.23 | 50 | 87.31 | 3.87 | 32 |
| 2013 | December | 12 | PA | 244.80 | 3.21 | 19 | 86.87 | 2.85 | 22 | 18.70 | 0.59 | 21 | 84.31 | 4.77 | 22 |
| 2014 | July | 21 | MA | 202.56 | 0.84 | 35 | 88.88 | 2.17 | 40 | 26.77 | 1.57 | 23 | - | - | - |
| 2014 | June | 26 | PA | 208.61 | 0.67 | 36 | 84.45 | 1.18 | 43 | 37.35 | 1.76 | 43 | 321.48 | 28.30 | 17 |
| 2014 | June | 25 | VA | 207.43 | 0.78 | 38 | 85.63 | 1.33 | 39 | 35.63 | 2.03 | 38 | 811.83 | 100.71 | 27 |
| 2014 | September | 9 | PA | 257.67 | 3.78 | 35 | 80.73 | 1.82 | 39 | 51.84 | 2.82 | 21 | 267.52 | 14.57 | 40 |
| 2014 | October | 15 | MA | 249.19 | 3.58 | 38 | 73.47 | 1.86 | 40 | 43.07 | 1.96 | 22 | | | |
| 2014 | October | 17 | PA | 247.93 | 3.63 | 37 | 67.92 | 1.17 | 46 | 30.38 | 2.06 | 40 | 130.31 | 4.08 | 17 |
| 2014 | October | 13 | VA | 247.94 | 3.78 | 33 | 71.65 | 1.96 | 37 | 48.95 | 2.29 | 18 | 95.29 | 4.68 | 33 |
| 2015 | July | 8 | PA | 153.58 | 0.95 | 36 | 72.99 | 1.27 | 40 | 57.01 | 3.24 | 18 | 254.00 | 10.81 | 40 |
| 2015 | October | 23 | PA | 190.49 | 3.63 | 28 | 63.92 | 2.32 | 29 | 43.38 | 2.80 | 29 | 124.62 | 14.91 | 6 |

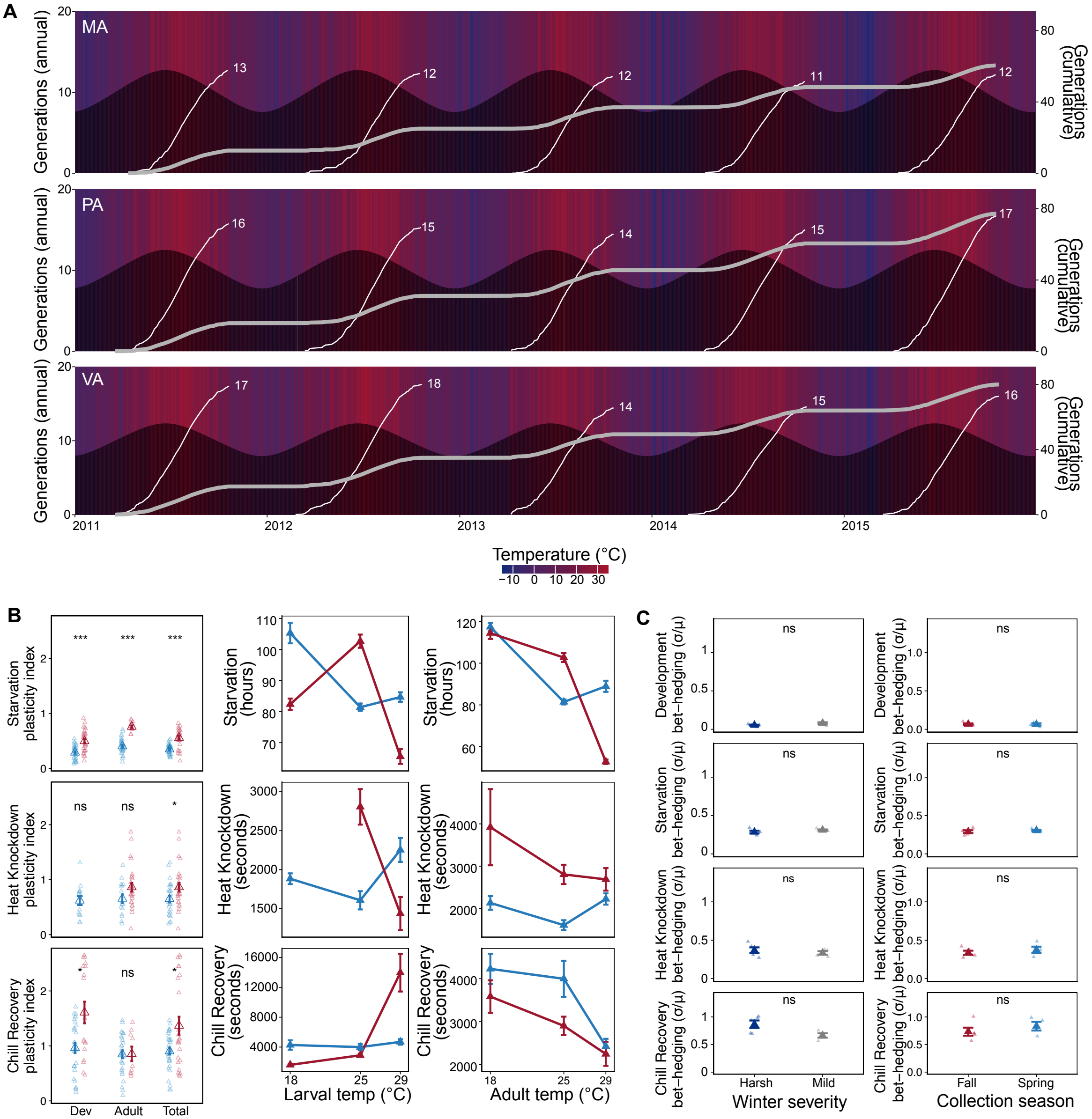

**fig. S1. Environmental context and phenotypic plasticity supplement.** (**A**) Principal component analysis biplot of meteorological variables for the Pennsylvania site, with sample trajectories colored by year. Table shows key winter (minimum temperature, days below 0°C, snowfall, wind ratio) and summer (maximum temperature, days above 32°C, precipitation) statistics for each year; H = harsh winter, M = mild winter. (**B**) Plasticity indices (left) and reaction norms for larval (middle) and adult (right) acclimation temperatures for spring (blue) and autumn (red) collections. Asterisks indicate significant seasonal differences in plasticity index (t-test, BH-corrected). (**C**) Bet-hedging indices (coefficient of variation of annual line means) separated by winter severity (left) and collection season (right) for each trait. Asterisks indicate significant differences (t-test, BH-corrected; *P<0.05, **P<0.01, ***P<0.001).

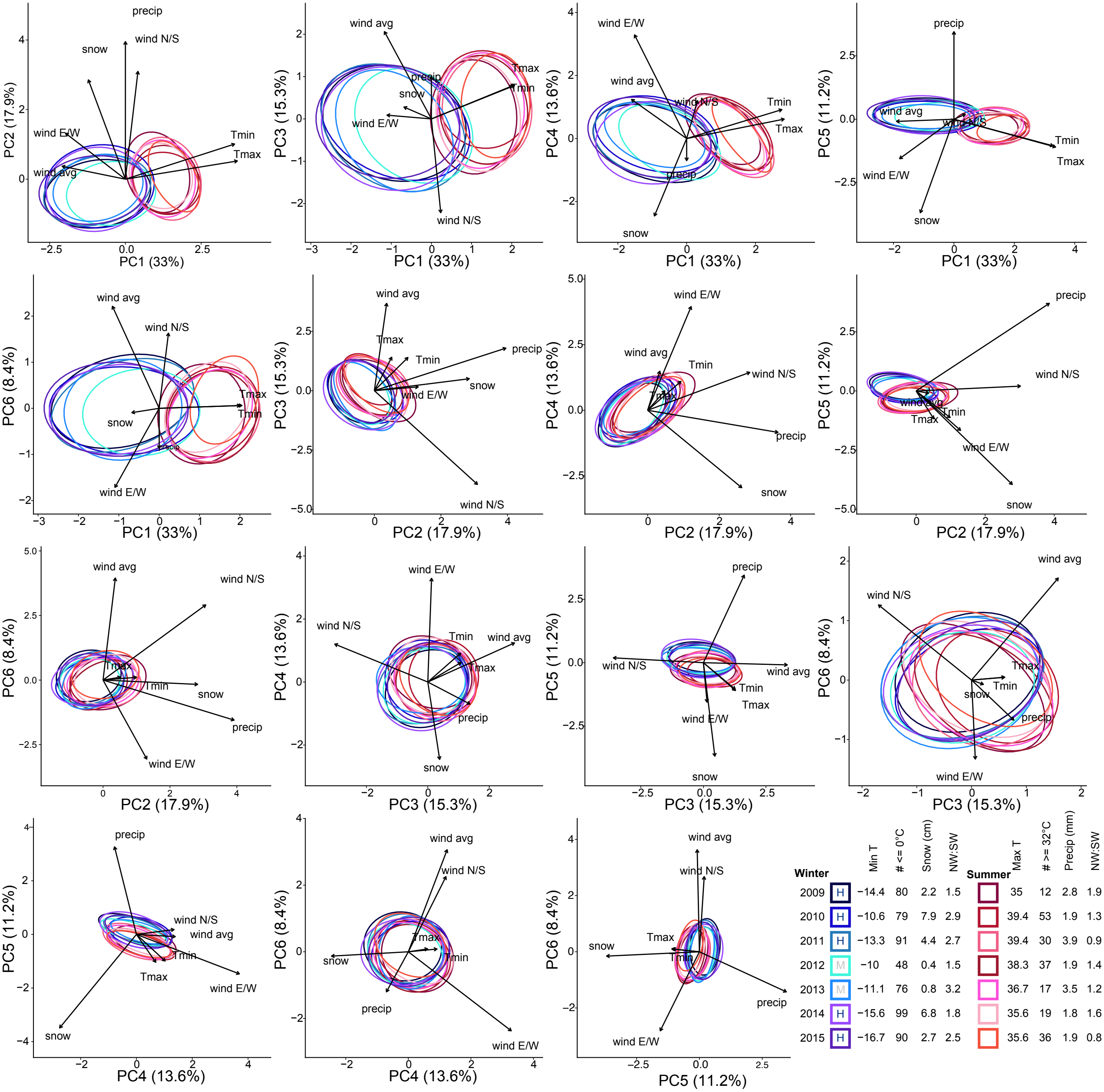

| Winter | | Min T | # <= 0°C | Snow (cm) | NW:SW | Summer | Max T | # >= 32°C | Precip (mm) | NW:SW |
|---|---|---|---|---|---|---|---|---|---|---|
| 2009 | H | −14.4 | 80 | 2.2 | 1.5 | | 35 | 12 | 2.8 | 1.9 |
| 2010 | H | −10.6 | 79 | 7.9 | 2.9 | | 39.4 | 53 | 1.9 | 1.3 |
| 2011 | H | −13.3 | 91 | 4.4 | 2.7 | | 39.4 | 30 | 3.9 | 0.9 |
| 2012 | M | −10 | 48 | 0.4 | 1.5 | | 38.3 | 37 | 1.9 | 1.4 |
| 2013 | M | −11.1 | 76 | 0.8 | 3.2 | | 36.7 | 17 | 3.5 | 1.2 |
| 2014 | H | −15.6 | 99 | 6.8 | 1.8 | | 35.6 | 19 | 1.8 | 1.6 |
| 2015 | H | −16.7 | 90 | 2.7 | 2.5 | | 35.6 | 36 | 1.9 | 0.8 |

**fig. S2. Environmental principal component analysis.** All pairwise principal component analysis biplots of meteorological variables for the Pennsylvania site, with sample trajectories colored by year. Table shows key winter (minimum temperature, days below 0°C, snowfall, wind ratio) and summer (maximum temperature, days above 32°C, precipitation) statistics for each year; H = harsh winter, M = mild winter.

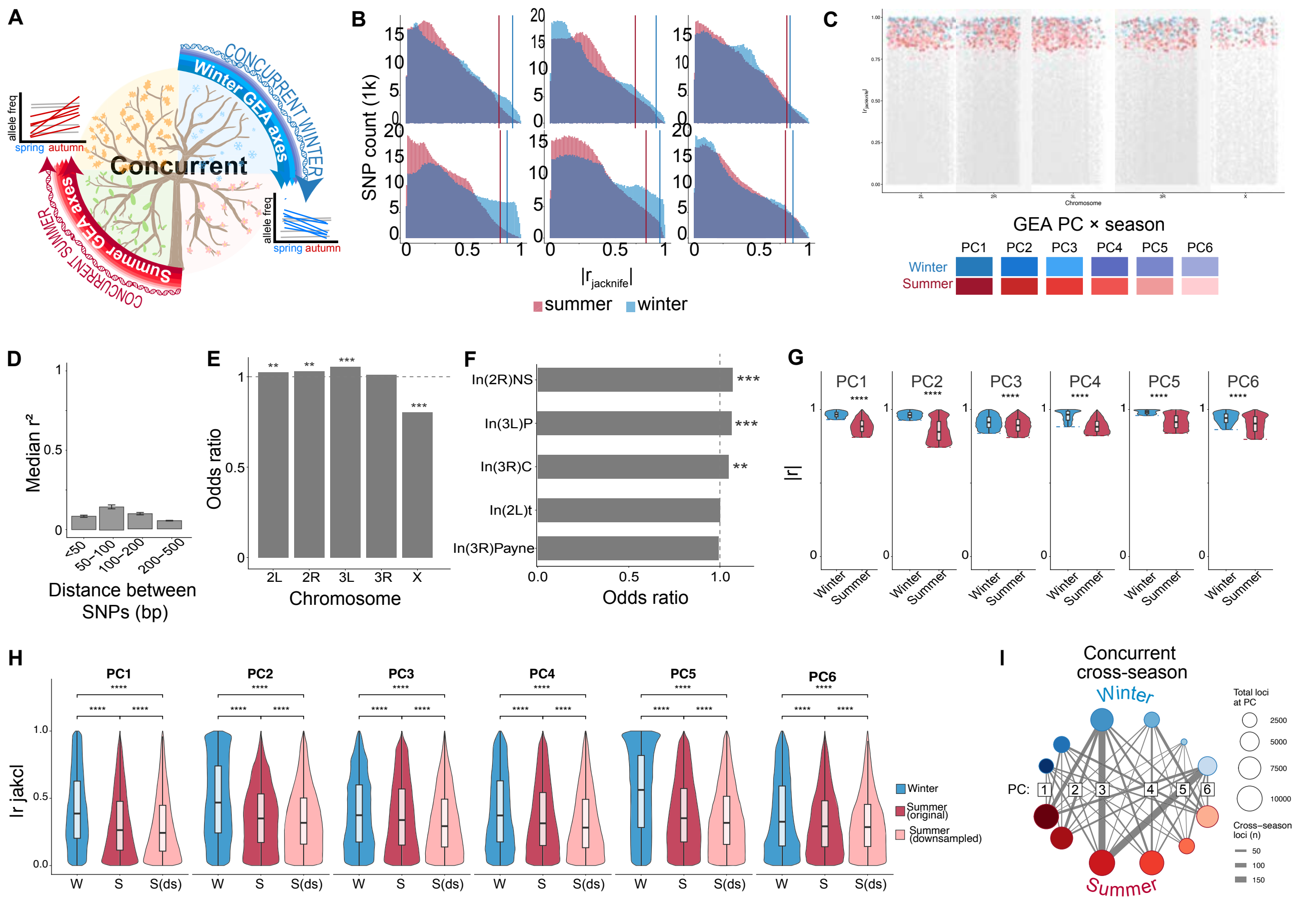

**fig. S3. GEA supplement: LD decay, empirical thresholds, chromosome and inversion enrichment, and per-PC correlation distributions.** (**A**) Schematic of the concurrent GEA approach: allele-frequency change is correlated against that season's environmental PC scores, tested separately for winter and summer. (**B**) Distribution of jackknife-corrected |r|, by PC and season (blue = winter, red = summer); vertical lines, each PC×season's permutation threshold. (**C**) Manhattan plot of jackknife |r| (gray, all SNPs), with SNPs significant in both seasons for the same PC highlighted. (**D**) Median r2 between significant concurrent GEA SNP pairs by physical distance (123 inbred lines); dashed line, LD-pruning threshold ($r^2$=0.5). (**E**) Chromosome enrichment of concurrent GEA SNPs (Fisher's exact test, BH-corrected; *P<0.05, **P<0.01, ***P<0.001); dashed line, obs/exp=1. (**F**) Cosmopolitan inversion enrichment of concurrent GEA SNPs (Fisher's exact test, BH-corrected); dashed line, odds ratio=1; ns, not significant. (**G**) Jackknife-corrected |r| for winter- and summer-associated SNPs, by PC (****P<0.0001). (**H**) Genome-wide jackknife |r| for winter (n=5), summer (n=6), and summer downsampled to n=5, by PC. (**I**) Routing network of SNPs significant for the same PC in both seasons; node size, total SNPs per PC; edge width, shared SNPs.

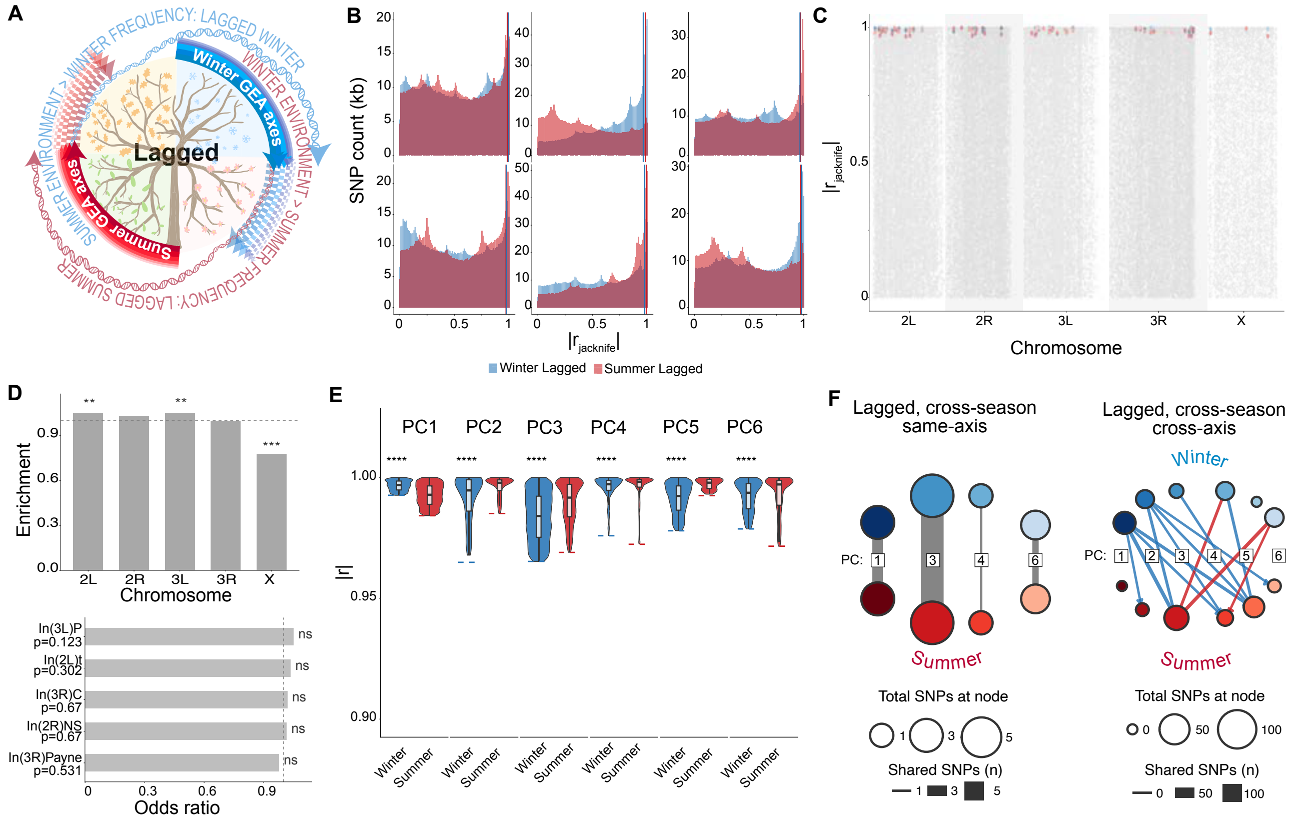

**fig. S4. GEA supplement for lagged loci: empirical thresholds, chromosome and inversion enrichment, and genomic position of shared loci.** (**A**) Schematic of the lagged GEA approach: winter allele-frequency change is correlated against the preceding summer's environmental PC scores, and summer change against the preceding winter's, tested separately for each direction. (**B**) Distribution of |r jackknife| across all tested SNP × PC × direction combinations, by PC; vertical lines, each PC × direction's empirical permutation threshold (winter-lagged, blue; summer-lagged, red). (**C**) Genomic position of the 59 loci significant in both lagged directions (colored by PC × direction), against the genome-wide distribution of jackknife |r| for all tested lagged loci (gray). (**D**) Chromosome enrichment (top) and cosmopolitan inversion enrichment (bottom) of significant lagged loci relative to the genome-wide background (Fisher's exact test, BH-corrected; **P<0.01, ***P<0.001); dashed lines, obs/exp = 1 and odds ratio = 1; ns, not significant. (**E**) Jackknife-corrected |r| for winter-lagged and summer-lagged SNPs, by PC (****P<0.0001). (**F**) Seasonal routing networks for lagged associations: loci shared between the same PC in opposite seasons (left) and between different PCs across seasons (right); edge width, shared loci; node size, locus count; edge direction reflects the greater share of loci for that PC pair.

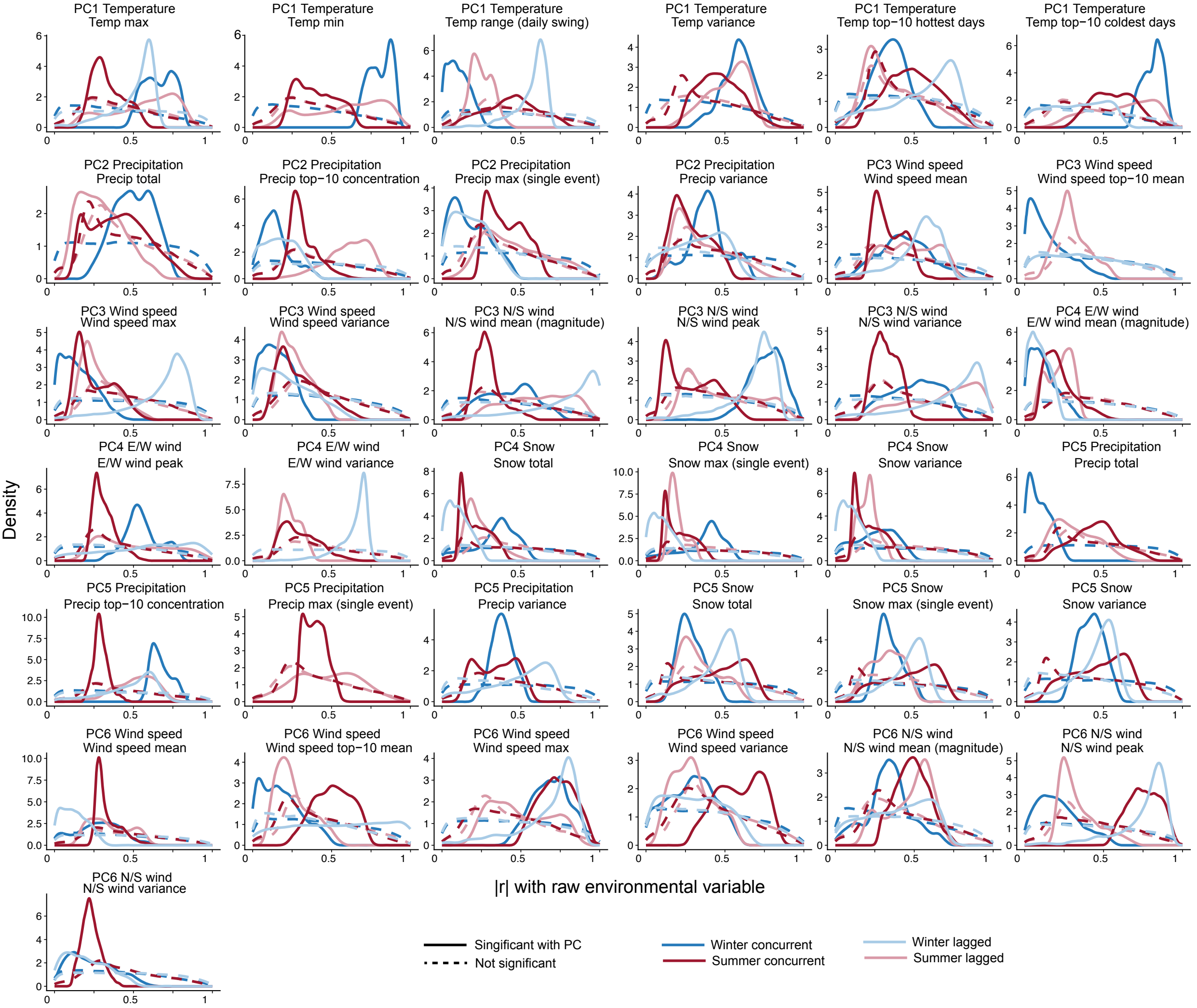

**fig. S5. Validation of PC-level associations against raw environmental variables.** Density distributions of |r| between each SNP and the raw environmental variable underlying a given PC, split by trigger-by-mode category (winter-concurrent, summer-concurrent, winter-lagged, summer-lagged) and by significance status (solid, significant; dashed, non-significant). Panels show only category × variable combinations passing Wilcoxon significance (BH-corrected $P<0.05$) between the significant and non-significant distributions. Across nearly every variable and category, significant SNPs show markedly higher |r| with the raw environmental measurement than non-significant SNPs, confirming that PC-level associations reflect genuine environmental signal rather than an artifact of the PCA itself.

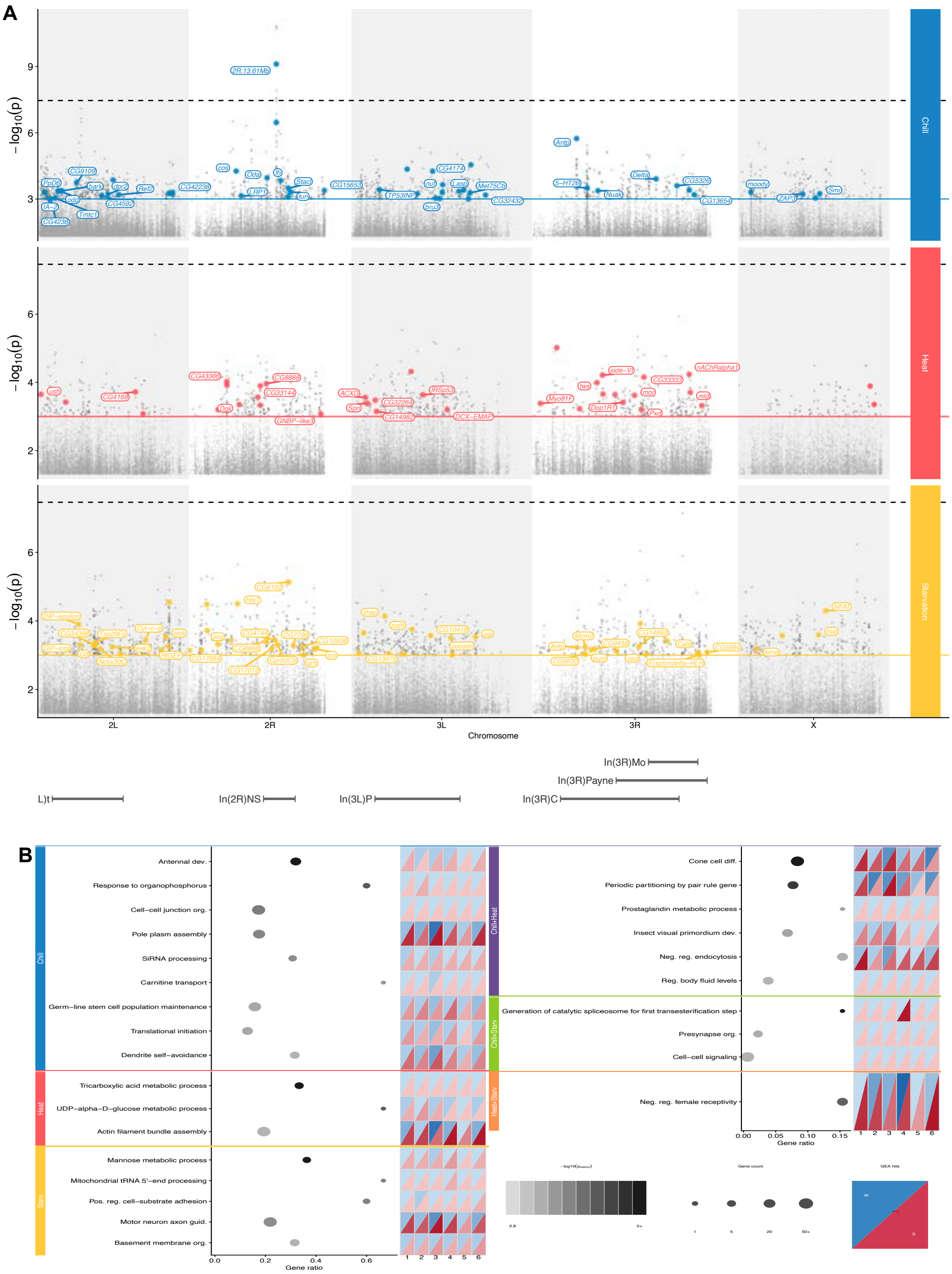

**fig. S6. GWAS–GEA convergence across trait-overlap categories.** (**A**) GWAS Manhattan plots (P<0.001) for chill, heat, and starvation (inbred PA lines). Gray = all SNPs; colored = also GEA-significant (concurrent, jackknife CI excl. zero). Labeled genes reach genome-wide significance. Dashed line = Bonferroni α=0.05; solid line = P=0.001. Inversions marked below. (**B**) GO enrichment (goseq, length-corrected, raw P<0.005, uncorrected) for six trait-overlap categories (singles left, pairs right). Dot size = gene count; color = −log10(P). Heatmaps show concurrent GEA hit counts by PC (1–6) and season (winter/summer triangles).

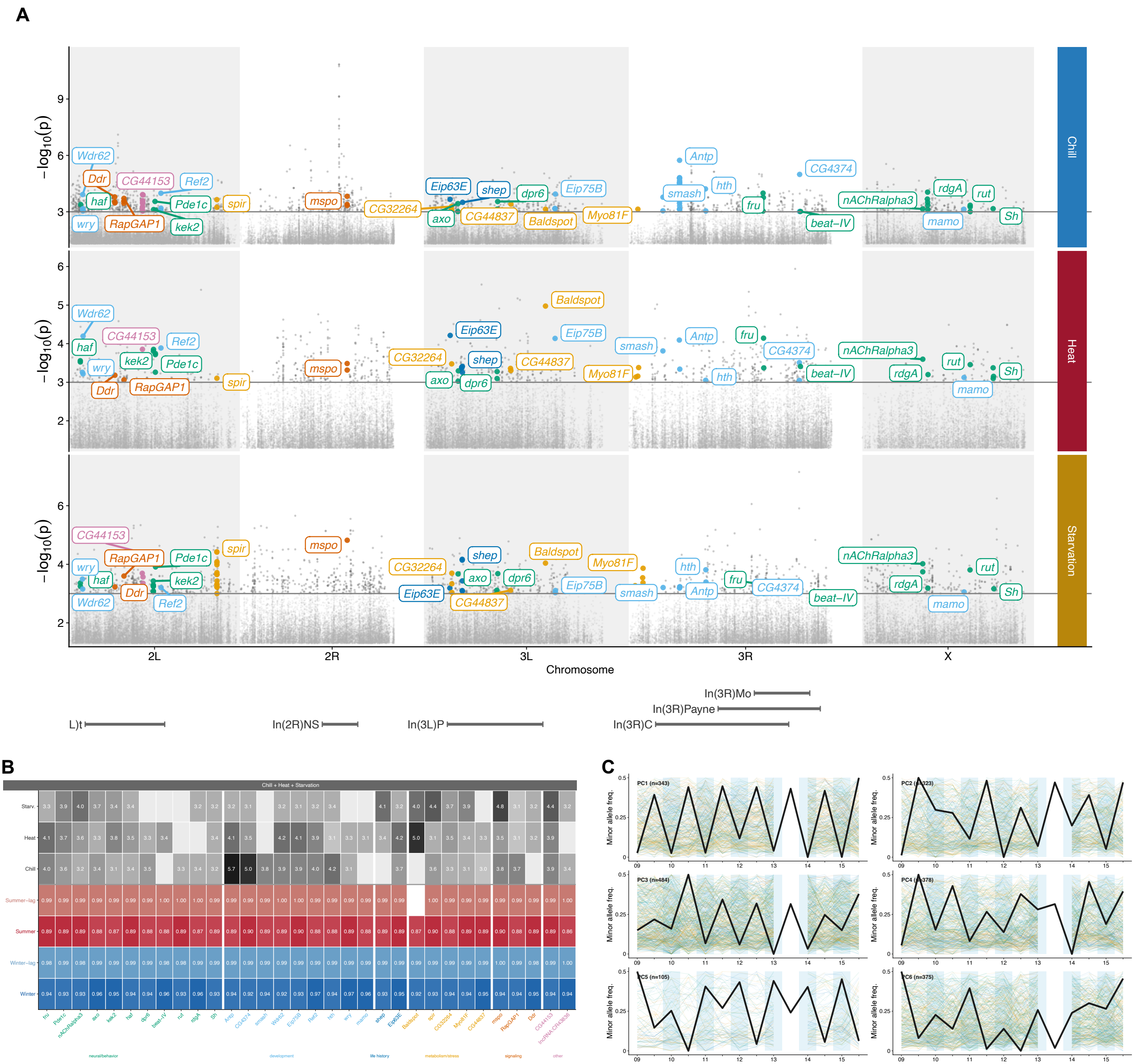

**fig. S7. Genomic and phenotypic architecture of the 32 triple-overlap genes.** (**A**) GWAS Manhattan plots (P<0.001) for chill, heat, and starvation, labeled by triple-overlap gene; label color = broad functional category (development, light blue; life history, dark blue; metabolism/stress, gold; signaling, orange; neural/behavior, green; other, pink). Gray = background SNPs. (**B**) GWAS effect (β; gray scale, darker = larger) and mean concurrent + lagged GEA |r| (blue = winter/winter-lag, red = summer/summer-lag; darker = concurrent, lighter = lagged) per gene. Inversions marked above. (**C**) Minor allele frequency trajectories (2009–2015) for loci significant on each environmental PC; thin colored lines = individual loci (colored by functional category), black = mean environmental PC score. Winter collections shaded blue.

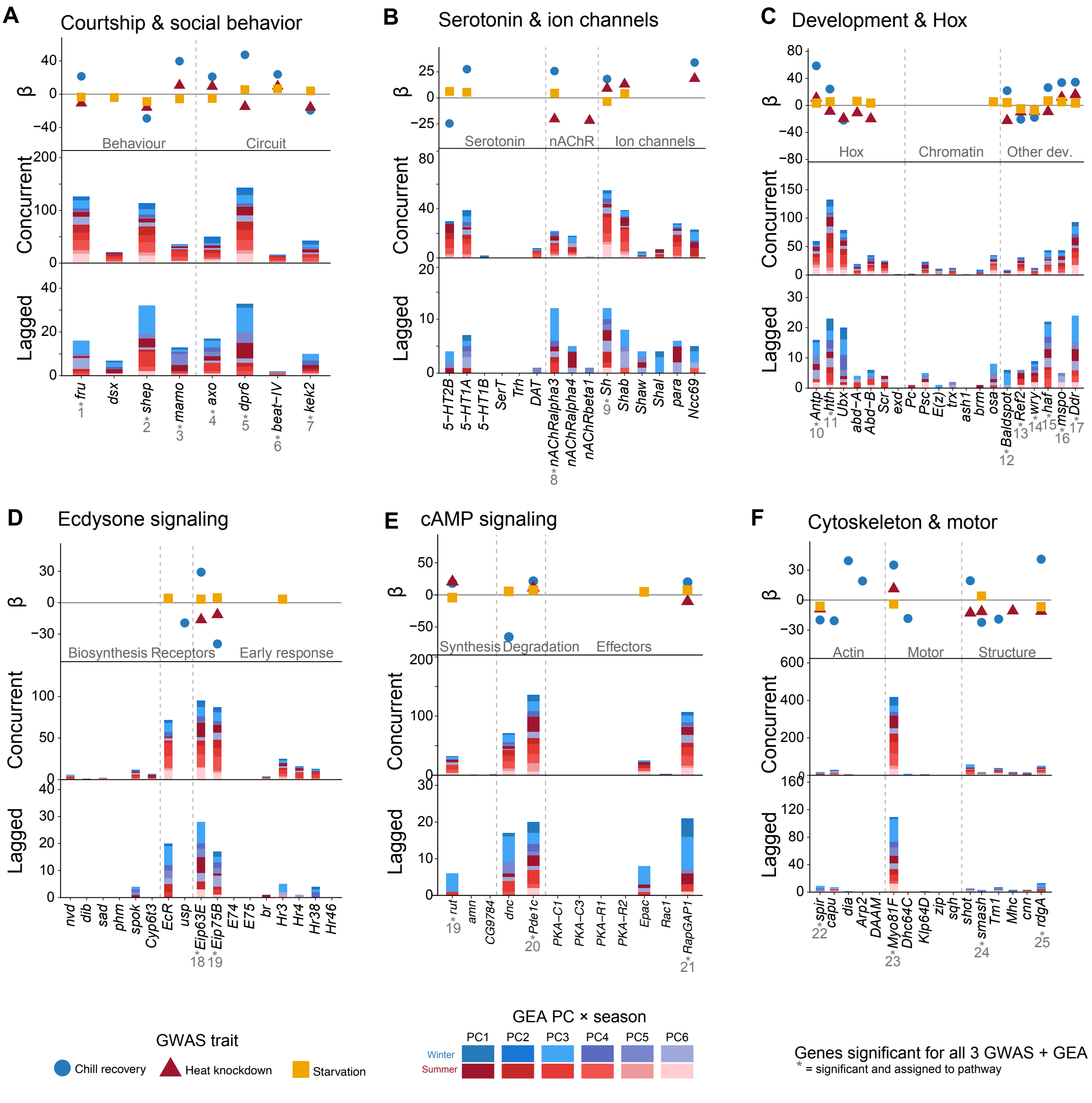

**fig. S8. GWAS for thermal and starvation tolerance identifies candidate loci overlapping extensively with seasonal GEA hits and clustering in functionally coherent pathways linking environmental axes to molecular mechanisms.** Genes containing both concurrent and lagged GEA loci (Shared) are enriched for GWAS hits relative to background genes, while genes with only concurrent or only lagged loci are significantly depleted (Fisher's exact test, BH-corrected; $^{**}P<0.01$, $^{***}P<0.001$). (A to F) Each pathway panel shows the GWAS effect size (β, genotype coefficient from a linear model of phenotype ~ genotype + season, upper) and GEA-significant SNP counts (lower, stacked by PC × season) for genes in six functional groups: (**A**) courtship and social behavior; (**B**) serotonin and ion channels; (**C**) development and Hox; (**D**) ecdysone signaling; (**E**) cAMP signaling; (**F**) cytoskeleton and motor proteins. β is shown for the most significant SNP per gene per trait. Genes are grouped into functional subcategories (dashed lines). GEA bar colors indicate contributing PC and season (blue shades, winter PCs 1–6; red shades, summer PCs 1–6). Asterisks denote genes among the 32 triple-overlap set. Six triple-overlap genes not assigned to pathways are: *Wdr62*, *CG44153*, *CG2264*, *CG44837*, *CG4374*, *lncRNA:CR43836*.

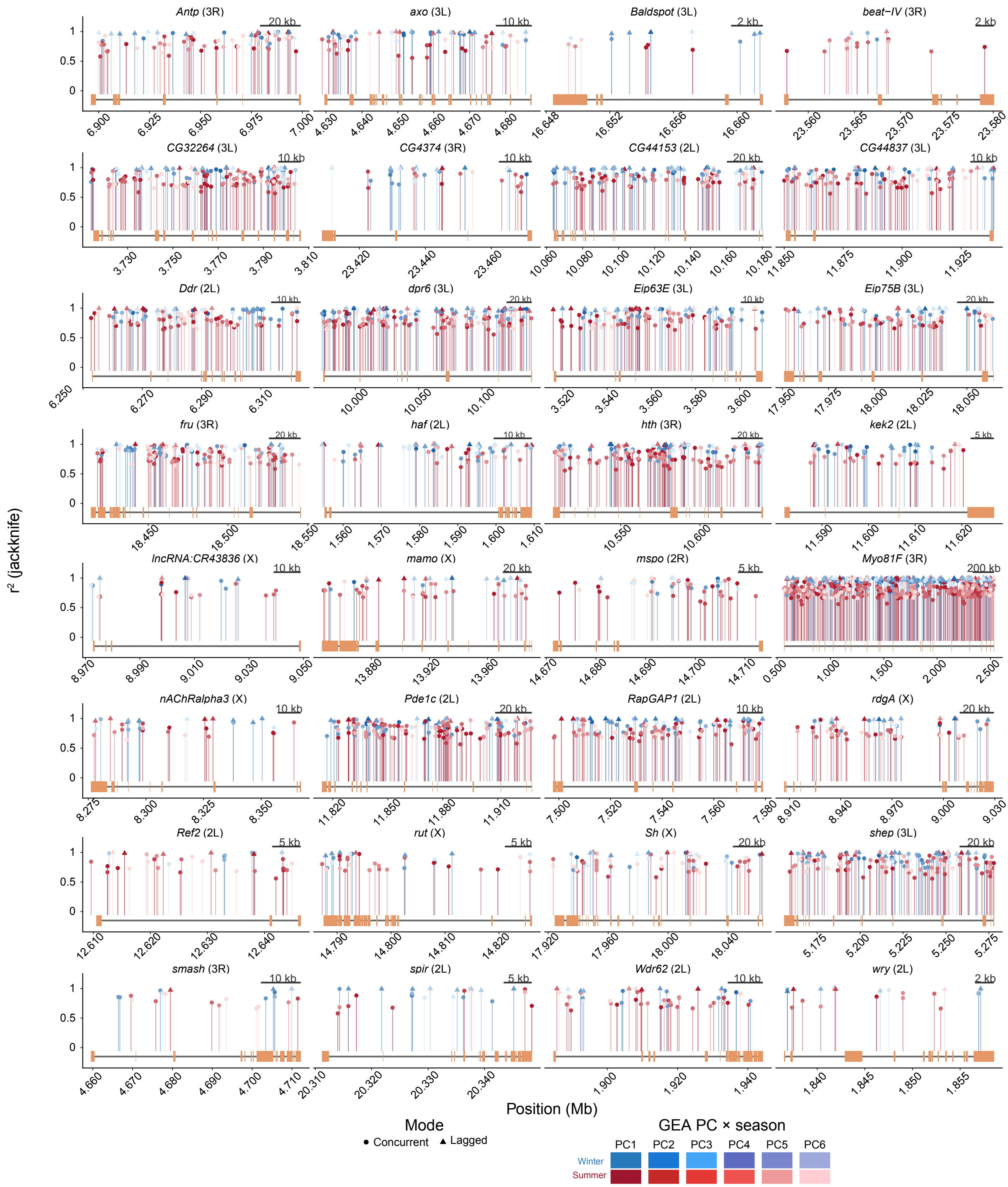

**fig. S9. Gene structure and GEA loci for the 32 triple-overlap genes.** Exon structure (orange) and intron backbone (gray line) for each gene, with significant concurrent (circle) and lagged (triangle) GEA loci plotted as lollipops (y = $r^2$ jackknife). Color = PC × season (blue shades, winter PC1–6; red shades, summer PC1–6; PC1 darkest). Chromosome noted per gene; scale bar length varies by gene (genes span 2 kb–2.5 Mb) and is individually labeled.

### References

48. V. Trotta, F. C. Calboli, M. Ziosi, D. Guerra, M. C. Pezzoli, J. R. David, S. Cavicchi, Thermal plasticity in Drosophila melanogaster: a comparison of geographic populations. *BMC Evol Biol* **6**, 67 (2006).

49. C. D. Schlichting, The Evolution of Phenotypic Plasticity in Plants. *Annual Review of Ecology and Systematics* **17**, 667–93 (1986).
